# A Joint Promoterome-Proteome Atlas Highlights the Molecular Diversity of Human Skeletal Muscles

**DOI:** 10.64898/2026.04.08.717200

**Authors:** Andrey Buyan, Guzel Gazizova, Viktor G. Zgoda, Nikita E. Vavilov, Nikita Gryzunov, Irina A. Eliseeva, Vladimir Nozdrin, Yulia Sergeeva, Angelina Titova, Leyla Shigapova, Alina V. Erina, Georgy Mescheryakov, Aysylu Murtazina, Ruslan Deviatiiarov, Alistair R.R. Forrest, Vsevolod Makeev, Yoshihide Hayashizaki, Daniil Popov, Elena Shagimardanova, Ivan V. Kulakovskiy, Oleg Gusev

## Abstract

More than 600 distinct skeletal muscles constitute up to 40% of the total mass of the human body. Human skeletal muscles differ in anatomical position, morphology, origin, and function, but the diversity of their molecular phenotypes, the gene expression and protein abundance profiles, remains poorly explored. Here, we report the large-scale CAGE-Seq promoterome profiling of 75 human skeletal muscles, complemented by 22 matched proteomes obtained with mass spectrometry. We identified 37001 transcribed regulatory elements and 1804 protein groups encompassing 1895 proteins, 80% of which demonstrated non-uniform expression across different muscles. The skeletal muscles of the eye, tongue, and diaphragm had the most distinctive molecular phenotypes, while the overall diversity was driven by hundreds of transcription factors with tissue-specific activity. By analyzing the allelic imbalance of CAGE-Seq reads, we discovered 6653 allele-specific single-nucleotide variants often coinciding with muscle-related GWAS SNPs, including muscle volume. Finally, we provide an interactive online atlas of transcriptomic and proteomic molecular phenotypes, facilitating further studies of gene regulation and heritable pathologies of skeletal muscles.

## Introduction

The human body contains more than 600 unique skeletal muscles that comprise approximately 40% of its total mass^1^. Besides morphological and positional differences, skeletal muscles vary in biochemical properties and molecular phenotypes, reflecting muscle etiology and functional needs. For instance, the expression of genes encoding sarcomere components, including myosin light and heavy chain isoforms, depends on the muscle origin, and is regulated by neural stimulation and concentrations of numerous steroid and thyroid hormones^2,3^. A complex interplay of external and internal stimuli results in diverse muscle composition, such as the varying fraction of slow- and fast-twitch fibers, and leads to different metabolic activity, contraction speed, stiffness, and even insulin sensitivity^4,5^. In the case of pathologies, these dissimilarities become critical as distinct muscle groups respond differently to myopathies caused by denervation, autoimmune and endocrine disorders, mitochondrial dysfunction, or muscular dystrophies^2,6–10^. In particular, the extraocular muscles are known to tolerate many forms of muscular dystrophies and aging, while having a distinct gene expression profile with a wide range of transcribed myosin heavy chain isoforms and genes involved in excitation-contraction coupling^11–13^.

Altogether, there is ample evidence of molecular heterogeneity of human skeletal muscles. However, systematic assessment of these differences requires a large number of samples taken from various parts of the body, including some that are harder to reach, like extraocular or deep skeletal muscles. Considering non-human vertebrates, there have been numerous studies of molecular phenotypes with RNA-Seq: transcriptomes of 47 skeletal muscles were profiled in pigs, 18 in horses, 11 in mice, and 10 in sheep, goat, and chicken^14–18^. These data highlighted an extraordinary diversity of skeletal muscle molecular phenotypes, with the intramuscular differences comparable to those between different body tissues, e.g., liver and cerebrum. In particular, genes encoding sarcomere components or proteins involved in energy metabolism, as well as HOX transcription factors, were the most variable across different skeletal muscles, reflecting different muscle fiber composition and origin.

In contrast to model or farm animals, human muscle transcriptomic or proteomic studies were severely limited by muscle availability, resulting in a bias towards biopsies taken from a few easily accessible sites, mainly located in the thigh muscles, with the maximum number of 6 different leg muscles whose transcriptomes were analyzed by Abbassi-Daloii T *et al.*^19–25^. In larger projects dedicated to exploring gene expression across human tissues and cell states, such as GTEx, Human Protein Atlas, or Human Cell Atlas, skeletal muscles are represented poorly, e.g., by a single limb muscle^21,26,27^; ENCODE data is marginally richer by a margin of only a few individual muscles (*gastrocnemius medialis*, *psoas*) and a few embryonic samples (from arm, leg, trunk and back)^28,29^. Finally, several integrative databases were constructed by integrating distinct datasets on human skeletal muscle gene expression: SKmDB, which contains multi-omic experiments for mainly embryonic tissues or *psoas* from the individuals of different age and focused on dynamic expression analysis^30^, MuscleAtlasExplorer, which integrates the results of 1654 microarray samples of 12 different muscles^31^, and MSdb, which combines the skeletal muscle RNA-Seq and scRNA-Seq experiments conducted mainly on healthy leg muscles before and after exercises or affected by diseases^32^. All in all, the overall molecular diversity of the skeletal muscles is explored only fragmentarily, and there is a clear need for a more comprehensive atlas encompassing different skeletal muscles from multiple body sites.

The gene expression diversity is not limited to tissue types, but also depends on individual genome variation, such as single-nucleotide variants (SNVs). In order to characterize the contribution of SNVs to muscle-related phenotypes, including muscle mass, hand grip strength, onset and severity of different muscle diseases, there have been numerous genome-wide association studies (GWAS)^33–42^. The majority of GWAS SNVs are located in non-coding regions, which heavily complicates the variant prioritization, assessment of causality, and validation. Despite successes in high-throughput techniques, the non-coding regulatory variants remain underexplored, and only a few of the muscle-related regulatory SNVs were validated experimentally^43^. Expanding the collection of annotated regulatory variants would enable better characterization of the genetic component of myopathies and other conditions in which no potentially causative protein-coding mutations have been observed. For skeletal muscles, several studies were already employing multi-omic analyses integrating DNA methylation, chromatin conformation, gene expression, and histone modification data, followed by identification of quantitative trait loci (QTLs) or allele-specific variants (ASVs)^44–46^. The latter approach seems specifically promising as, profiting from the intra-individual genetic variants and comparing the allelic counts at heterozygous SNV, it reduces confounding effects by inherent inter-individual normalization and allows variant prioritization even when the sample size is relatively small^47^.

Following the trail established by the well-known FANTOM (functional annotation of mammalian genome) project^48^, here we present FANTOMUS, Functional ANnoTation Of human skeletal MUScle genome (https://fantomus.autosome.org), a skeletal muscle-centric atlas of transcribed regulatory elements (TREs) accompanied by protein abundance profiles. For FANTOMUS, we performed Cap Analysis of Gene Expression (CAGE-Seq) of 222 samples from 75 skeletal muscles, as well as Liquid Chromatography - tandem Mass Spectrometry (LC-MS/MS) of 60 samples from 22 muscles, and identified 37 thousand active TREs and 1804 protein groups (encompassing 1895 individual proteins, indistinguishable within each group due to a limited number of detected peptides). These data were used to estimate the intermuscular gene expression heterogeneity at promoter and protein levels, revealing the expression of 80% of genes and proteins being non-uniformly distributed across muscles, with the most distinct expression profiles of the extraocular muscles, tongue, and diaphragm. Motif activity response analysis confirmed extreme molecular diversity of human skeletal muscles with hundreds of transcription factors demonstrating differential activity. On top of that, CAGE-Seq allele-specific analysis identified 6,5 thousand single-nucleotide variants (SNVs) with the imbalanced transcriptional signal between alleles. These regulatory promoter variants were enriched for muscle-related GWAS traits, such as muscle volume, and selected case studies were validated using a luciferase reporter assay.

Taken together, here we present FANTOMUS as the largest to date unified atlas of human skeletal muscle expression profiles, providing promoter and enhancer activity, protein abundance, and the effects of the individual variants on muscle transcriptional profile, and thus enabling the in-depth analysis of intermuscular and interindividual molecular heterogeneity.

## Results & discussion

### Transcriptomic and proteomic atlas of human skeletal muscles

To identify genomic locations and quantitatively estimate the activity of transcribed regulatory elements (TREs, promoters, and transcribed enhancers) in different skeletal muscles, we performed CAGE-Seq of 222 autopsy samples from 75 distinct muscles of 4 individuals (3 males and 1 female, 56-67 years old), followed by skeletal muscles grouping based on their affiliation with different parts of the body (**Figure 1a,b, Supplementary Figure 1a, Supplementary Data 1**). CAGE-Seq bioinformatics processing (quality control, peak calling, and filtering, see **Methods** and **Supplementary Figures 1b-d and 2**) yielded 37 thousand reliable TREs, including unidirectional promoters of 18329 genes and 881 bidirectionally transcribed enhancers^49^ (**Supplementary Data 2**). An illustrative example shown in **Figure 1c** displays the fast myosin heavy chain (MYH) gene cluster encompassing several common (*MYH1*, *MYH2*), fetal (*MYH3*, *MYH8*), and eye muscle-specific (*MYH4*, *MYH13*) genes, as well as a *linc-MYH* (ENST00000585303) marking the recently discovered super-enhancer^50^.

**Figure 1.**
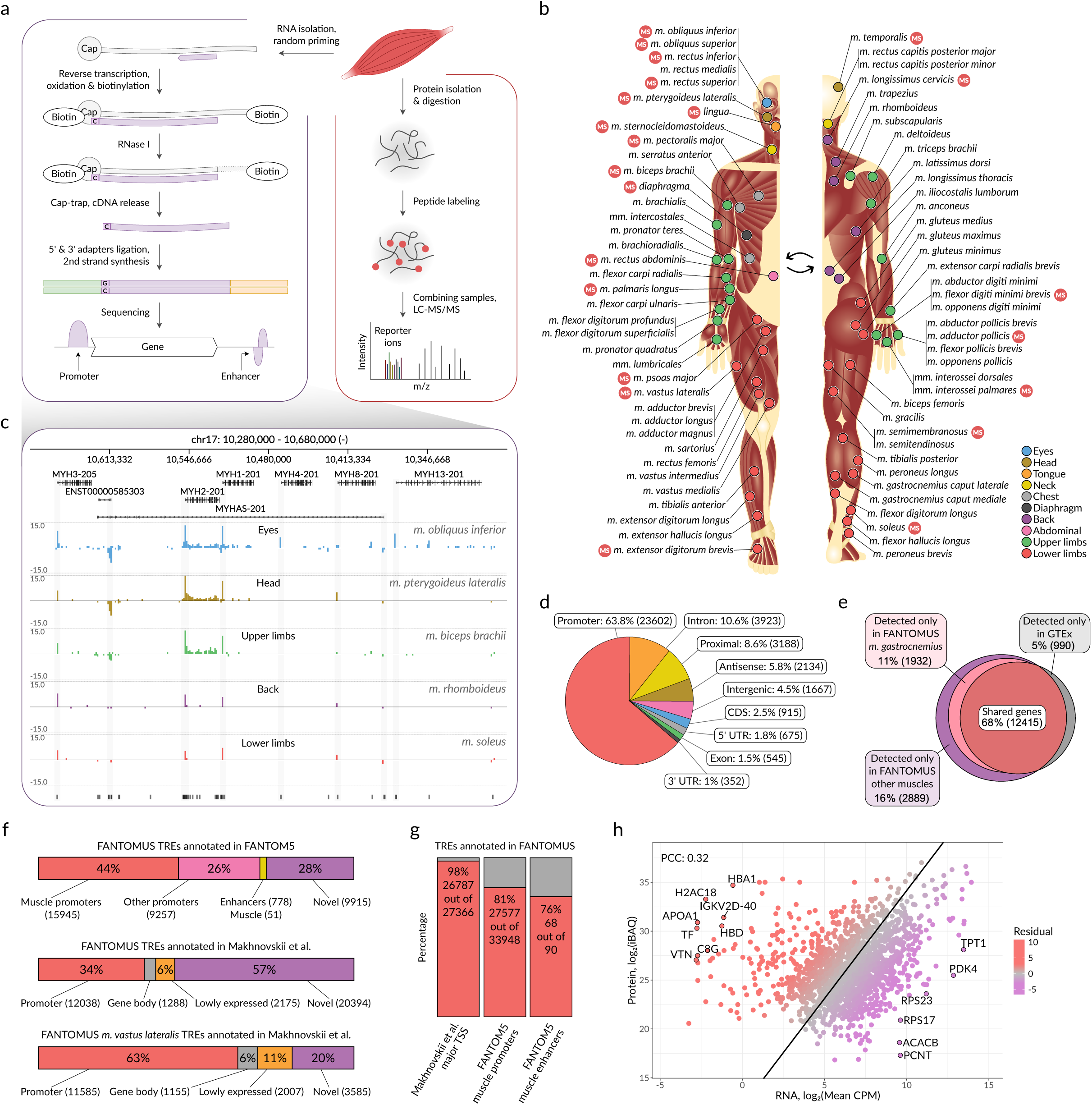
Transcriptomic (CAGE-Seq) and proteomic (MS) molecular phenotypes of 75 human skeletal muscles. **a**, Schematic representation of CAGE-Seq (left) and mass spectrometry (right). **b**, Anatomical location of skeletal muscle samples along the human body. CAGE-Seq was performed for 75 muscles; mass spectrometry was performed for a representative subset of 22 muscles (marked as MS). Colors denote body parts. **c**, Genomic view of the fast MYH gene cluster with the CAGE-Seq profiles for 5 skeletal muscles. Y-axis: the number of reads reflecting the capped 5’ transcript ends, log_2_, bottom (negative): direct strand, top (positive): reverse complementary strand. The lowest track denotes individual TREs (black bars); the major promoter of each gene is highlighted in grey. **d**, Categorical distribution of TREs across genomic regions. **e**, Protein-coding and lncRNA genes detected in FANTOMUS and GTEx skeletal muscle samples. **f**, Percentage of FANTOMUS TREs detected in FANTOM5 (top) and Makhnovskii *et al.* (middle); percentage of FANTOMUS *m. vastus lateralis* TREs detected in Makhnovskii *et al.* (bottom). **g**, Percentage of TREs from different datasets detected in FANTOMUS. **h**, Correlation between the RNA (X-axis, mean log_2_CPM) and protein (Y-axis, log_2_iBAQ) abundance. Overrepresented proteins in MS (red) and transcripts in CAGE-Seq (purple) are highlighted, and the extreme outliers are labeled; the solid line represents Deming regression, color scale reflects the residuals. Only autosomal genes and TREs were considered in **(e)**, **(f)**, **(g),** and **(h)**. LC-MS/MS: liquid chromatography-tandem mass spectrometry, CAGE-Seq: cap analysis of gene expression, TRE: transcribed regulatory element, iBAQ: intensity-based absolute quantification value, CPM: counts per million, PCC: Pearson correlation coefficient, CDS: coding sequence, UTR: untranslated region.

**Figure 2.**
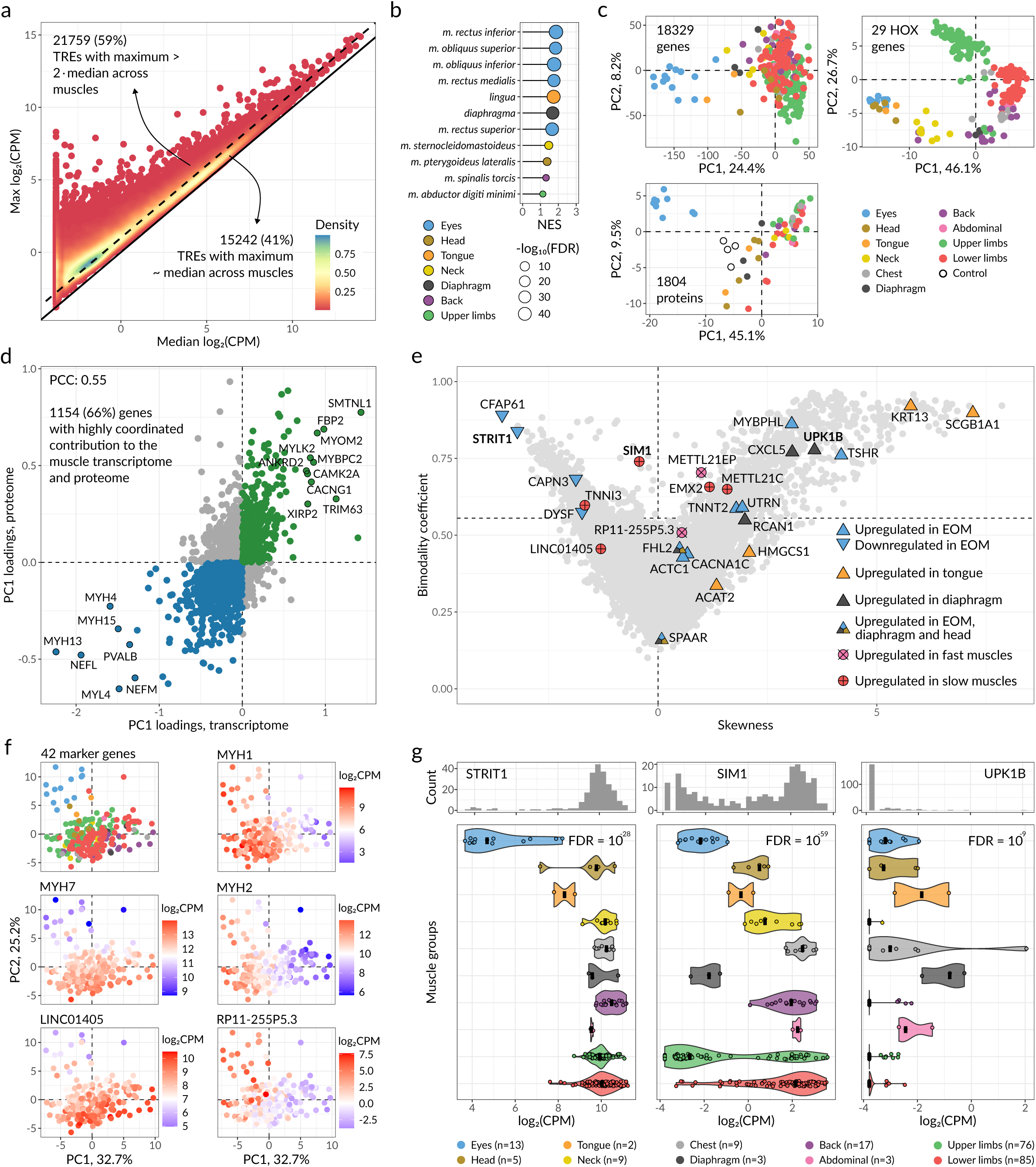
Gene expression and protein abundance heterogeneity across skeletal muscles. **a**, TRE activity across muscles: the maximal log_2_CPM (X-axis) plotted against the median (Y-axis). Diagonal: y = x, dashed line: max CPM > 2x median, color scale reflects the points density. **b**, Skeletal muscles characterized by the most unique transcriptional profiles, only 11 muscles significantly enriched (FDR < 0.05) by uniquely expressed TREs are shown. X-axis: GSEA NES, color: muscle group, point size: -log_10_FDR. **c**, PCA of the 222 muscle samples based on gene-level log_2_CPM values (top left) or a subset of 29 HOX genes (top right), and 60 muscle samples with 4 pooled controls based on protein-level log_2_Intensity values (bottom left). Axes: the first two components; color: muscle group. **d**, PC1 loadings from transcriptome- (X-axis) and proteome-based (Y-axis) PCA from **(c)** for common autosomal genes. Color denotes genes consistently up- (blue) or downregulated (green) in EOM, other non-somitic muscles, tongue and diaphragm. Extreme outliers discussed in the text are additionally labeled. **e**, Scatter plot demonstrating the skewness (X-axis) and bimodality coefficient (Y-axis) for 16033 genes significantly differentially expressed across muscles, ANOVA FDR < 0.05. Individual examples discussed in the text are highlighted and labeled; shape and color denote the expression pattern. **f**, PCA based on the gene-level log_2_CPM values of 42 fast- and slow-twitch marker genes from Abbassi-Daloii *et al.*^25^. Axes: the first two components; color: muscle group (top left panel) as in **(c)** or log_2_CPM values of individual genes. **g**, Examples of the left-skewed (left), bimodal (middle), and right-skewed (right) gene expression distributions of *STRIT1*, *SIM1*, and *UPK1B*, respectively. Histograms for gene-level log_2_CPM values (top) and violin plots for separate muscle groups (bottom), FDR-corrected ANOVA *P*-values are labeled at the top, color denotes muscle groups, bars represent the medians. TRE: transcribed regulatory element, CPM: counts per million, FDR: false discovery rate, GSEA: gene set enrichment analysis, NES: normalized enrichment score, PC: principal component, PCC: Pearson correlation coefficient, EOM: extraocular muscles.

The majority of the identified TREs are located in the promoter regions of genes annotated in MANE (v.1.1) and GENCODE (v.44), comprising 63.8% of the total number of clusters (**Figure 1d**). Focusing on the subset of bidirectionally transcribed enhancers^49^, they were preferably located within intronic, proximal, and intergenic regions, and enriched with enhancer-specific H3K4me1 histone modification according to the Roadmap Epigenomics^51^ and ENCODE^52^ data, and with H3K18 lactylation mark of active tissue-specific enhancers^53^ (**Supplementary Figure 1e-f**).

Next, we compared FANTOMUS with GTEx (v.10)^54^ RNA-Seq data on the gene level. Overall, 72% of genes detected in FANTOMUS were also detected in the GTEx skeletal muscle tissue transcriptome, increasing to 86% when considering only a subset of FANTOMUS genes expressed in the GTEx-matched *m. gastrocnemius* (**Figure 1e**). Quantitative comparison of the expression estimates between FANTOMUS (summing up read counts for all unidirectional TREs of a gene) and GTEx also displayed the agreement between CAGE- and RNA-Seq data, with the Pearson correlation coefficient (PCC) of 0.86 (**Supplementary Figure 3a**).

Next, we performed the TRE-level comparison between FANTOMUS, FANTOM5 promoters and enhancers^48^, and transcription start site (TSS) clusters from Makhnovskii *et al.*^55^. All three datasets are the results of CAGE-Seq; FANTOM5 covers several skeletal muscles, including extraocular muscles (EOM), while Makhnovskii *et al.* data came from *m. vastus lateralis* biopsy samples.

Annotating FANTOMUS with FANTOM5 data, the majority of the identified TREs are indeed active promoters and enhancers (**Figure 1f**); notably, the eye-specific FANTOM5 promoters were upregulated in FANTOMUS EOM samples (**Supplementary Figure 3b**). Further, we also detected nearly ten thousand reliable regulatory elements not presented in FANTOM5 data but passing the strict quality control in FANTOMUS (see **Methods** and **Supplementary Figures 3c**). Considering the newer data from Makhnovskii *et al.*, it included only 34% of FANTOMUS TREs as active promoters, but this value increased to 63% when considering a subset of FANTOMUS TREs active in *m. vastus lateralis* (**Figure 1f**).

We also evaluated the FANTOMUS sensitivity by estimating the percentage of detected major TSS clusters from Makhnovskii *et al.*, as well as muscle promoters and enhancers from FANTOM5, which resulted in 98%, 81%, and 76% of them, respectively, being successfully captured in FANTOMUS (**Figure 1g**).

To complement the promoter atlas at the protein level, we performed tandem mass spectrometry (MS) for a representative subset of 60 samples (**Supplementary Data 1**) and estimated the relative abundance for 1804 protein groups with at least 2 detected peptides for 22 skeletal muscles (**Figure 1a,b, Supplementary Figure 3e, Supplementary Data 2**). As expected, MS preferably captured highly abundant proteins, including sarcomere components, ribosomal proteins, and translation machinery components, with the tropomyosin alpha-3, alpha-1, and beta chains, actin alpha 1, and myosin light and heavy chains being the most abundant (**Supplementary Figure 3f**).

The total RNA level (measured as mean counts per million, CPM) and protein abundance (intensity-Based Absolute Quantification value, iBAQ) of 1750 autosomal genes detected both with MS and CAGE-Seq yielded a PCC of 0.32 (**Figure 1h**). We then focused on the outlier genes with either over- or underabundant protein levels. The former (overabundant proteins) were mostly enriched by extracellular matrix, immune, and plasma proteins, which likely reflect samples infiltration with blood, as well as stable nucleosome components, while the latter were represented by ribosome and proteasome components as well as proteins involved in cell cycle regulation, presumably reflecting strict translational control of these genes limiting the protein output despite mRNA availability^56,57^ (**Supplementary Figure 3g**). Besides the general correlation, we estimated sample-level correlations between normalized CAGE-Seq CPM values and intensities from MS, **Supplementary Figure 3h**. Here, EOM showed the highest median Spearman correlation coefficient (SCC) of 0.34 (compared to 0.15-0.24 of other muscle groups), suggesting that the post-transcriptional layer of gene expression regulation in these tissues is much weaker than in the rest of the skeletal muscles.

### Skeletal muscles have diverse molecular phenotypes

#### Activity profiles of transcribed regulatory elements highlight unique molecular phenotypes

TRE activity estimates across 75 muscles enable classifying individual TREs into "house-keeping" (stably expressed in the majority of the skeletal muscles) or "specific to particular muscle(s)" by comparing their maximal versus median expression (log_2_CPM) across all samples (**Figure 2a** and **Supplementary Data 3**). Of them, 21759 (59%) had a maximum expression value no less than twice the median and were indeed depleted of house-keeping promoters according to FANTOM5^48^: 1089 (5%) against 6192 (41%) from 15242 remaining TREs. By assigning each TRE to the muscle where it showed the maximal expression and ordering TREs by the difference between the maximal and median expression, we performed ranking analysis (as in the gene set enrichment analysis, GSEA) and identified muscles with the most uniquely expressed promoters and enhancers (**Figure 2b**).

As muscles from three groups, EOM (extraocular muscles), tongue, and diaphragm, showed the most significant enrichment with tissue-specific TREs (FDR < 10^-33^), we then compared them directly to the rest of the samples in terms of the differential gene expression and protein abundance, **Supplementary Data 4**. As a result, we identified 9302, 2161, and 2743 genes differentially expressed at FDR < 0.05 and |log_2_FC| > 0.5 in EOM, tongue, and diaphragm, respectively, as well as 412, 25, and 11 differentially abundant protein groups. The eye-specific genes were enriched by the processes of neurogenesis and retina homeostasis^2,11^, while on the protein-level, EOM were enriched by aerobic respiration, consistent with a higher level of oxidative metabolism in these muscles and their increased susceptibility to mitochondrial diseases^58–60^ (**Supplementary Figure 4a**). Tongue uniquely expressed genes involved in epidermis development and humoral immune response, and contained higher levels of cell adhesion proteins. Finally, diaphragm demonstrated a higher expression level of immune-related genes, including phagocytosis and neutrophil chemotaxis, which were shown to promote muscle regeneration^61^, as well as an increased level of proteins involved in oxidative phosphorylation, similar to EOM.

#### Assessing muscle RNA- and protein-level heterogeneity

Next, we performed the principal component analyses of muscle samples using the expression estimates for 18329 genes and the abundances of 1804 protein groups (**Figure 2c**), which segregated the samples similarly: the first principal components (PC1) of both transcriptomic and proteomic data distinguished EOM, tongue, diaphragm, neck, and head from the other muscles explaining 24.4% (transcriptome) and 45.1% (proteome) of the total variance. This layout likely reflects the unique non-somitic origin of craniofacial muscles, which are known to express a special repertoire of sarcomeric genes^2^. Indeed, the top genes and proteins contributing to PC1 were enriched by gene ontology terms related to embryonic morphogenesis, muscle development and sarcomere organization, including such genes as myosin heavy and light chains (e.g., extraocular *MYH4*, *MYH13* and *MYH15*, slow-twitch *MYH7*, *MYL2* and *MYL3*), regulators of craniofacial development and neurogenesis (*ALX4*, *CTXN3*, *SPOCK3*, *NXPH2*, and *LMX1A*), satellite cells and myoblasts differentiation (*LRTM1*, *BARX2* and *CSRP3*), see **Supplementary Figure 4b,c**.

The designation of the second principal component depended on modality: the transcriptome distinguished upper and lower limbs based on genes involved in embryonic development (e.g., *HOXC10, HOXC9*, and *HOXC6*) and lipid metabolism (*HMGCS2, MSS51*, and *LIPG*), while the proteome separated muscles based on the reactive oxygen species-related and inflammatory proteins (encoded by *SAA1*, *MRC1*, *LYZ*, *CRP*, and *C1S*). Notably, the expression profile of HOX genes in adult skeletal muscle resembled that of embryonic development and was distinct for individual body parts, in agreement with the findings reported for non-human animals^14,16^, see **Figure 2c** and **Supplementary Figure 4d**.

We then directly compared the contribution to the first principal component of 1750 autosomal genes detected in both transcriptome and proteome. A group of 1154 genes with consistent signs in the proteome and transcriptome PC1 loadings separated diaphragm, tongue, extraocular, head, and neck muscles from those of other somitic muscle groups (**Figure 2d**). The group comprised EOM-specific MYH and neurofilaments, most likely reflecting the EOM-specific innervation^11^, and parvalbumin, which is known to serve as the Ca^2+^ buffer in murine fast muscle but has significantly lower expression in humans^62^. The expression of genes associated with glycolytic phenotype (namely *SMTNL1*^63^ and *FBP2*^64^), as well as fast skeletal myosin-binding protein-C and M-protein, showed the opposite trend and was decreased in EOM. Eye, head, and tongue muscles also showed downregulation of *XIRP1-2*, *ANKRD2*, and *CCN1*, see **Supplementary Figure 5a**, suggesting a reduced response of these muscles to stretch and mechanical strain^65–67^.

Finally, we detected a consistent decrease in muscle atrophy-associated gene expression, namely *TRIM63*, *FBXO32*^68^, *CACNG1*^69^*, FOXO3, FBXO21*^70^, and *PHAF1*^71^, in EOM, tongue, and diaphragm, indicating the reduced susceptibility of these muscle groups to wasting, e.g., with reduced muscle activity or aging, see **Supplementary Figure 5b**.

We then specifically explored the proteasome core and regulatory subunit gene expression, as related to the ubiquitin-proteasome system involved in muscle atrophy^72^, in terms of the overall contribution to PC1. The core subunit genes were downregulated in EOM, head, tongue, and diaphragm, while only those encoding the immuno-subunits (*PSMB8*), PA28 subunits (*PSME1* and *PSME2*), and proteasome inhibitor (*PSMD5*) were upregulated, possibly due to the increased endogenous IFNγ stimulation, which was shown to improve muscle regeneration^73–75^ (**Supplementary Figure 4e**). In addition, we found a generally increased protein abundance of complex I subunits in EOM, suggesting an increased level of oxidative metabolism (**Supplementary Figure 4e**).

The ribosomal proteins were recently reported among the main contributors to the heterogeneity of individual muscle fibers^76^, yet we did not detect any significant variation of gene expression and protein abundance across muscles. According to our data, the major differentially expressed ribosomal genes at both RNA and protein levels were *RPL3* and its muscle-specific paralog *RPL3L*^77^, which demonstrated the opposite tendencies being up- and downregulated in EOM, tongue, and diaphragm, respectively, see **Supplementary Figure 4e**. *RPL3L* was shown to be downregulated under the hypertrophic stimulus^78^ and Duchenne muscular dystrophy (DMD), presumably as a compensatory mechanism, as its knockdown generally enhanced muscle function^79^. Finally, mitochondrial ribosomal proteins were upregulated in EOM, likely reflecting a higher level of oxidative metabolism and mitochondrial content.

#### The majority of genes and proteins are differentially expressed across skeletal muscles

ANOVA across all 75 and 22 muscles revealed 16033 genes (87%) and 1469 proteins (81%) passing FDR < 0.05, respectively, i.e., having a significant differential expression. Of those, 3908 (21%) and 522 (29%) had highly skewed (absolute skewness over 1) or bimodal (bimodality coefficient^80^ > 5/9) expression patterns across muscles, see **Figure 2e**, **Supplementary Figure 4f**, and **Supplementary Data 3**.

The group with skewed expression included genes known to be down- or upregulated in specific muscles, like age-related marker *CFAP61*^19^ and SERCA activity enhancer *STRIT1*^81^, which are downregulated in EOM, or *UPK1B* upregulated in diaphragm, in agreement with the previous observations^16^ (**Figure 2g**). We further detected highly skewed genes preferably expressed in one of the three unique muscle groups described earlier, namely EOM, tongue, and diaphragm. The former included thyroid-stimulating hormone receptor and cardiac isoforms of genes involved in excitation-contraction coupling (*CACNA1C*) and sarcomere functioning (*ACTC1, TNNT2, MYBPHL*), which are known to be upregulated in EOM^13,82,83^. Keratins, uteroglobin, and cholesterol biosynthesis genes, namely *ACAT2* and *HMGCS1*, were found to be specifically expressed in the tongue, while *CXCL5* and *RCAN1* demonstrated diaphragm-specific expression, possibly reflecting its high vascularization associated with an increased immune cell recruitment^61^ and oxidative stress^84^. *FHL2* and *SPAAR* are involved in muscle development and regeneration^85–87^, and were upregulated in both EOM and diaphragm, as well as head muscles (**Supplementary Figure 5c**).

The group with bimodal expression primarily included genes differentially expressed between certain muscle groups, e.g., *HOXC10* upregulated in the lower half of the body or *SIM1* and *EMX2*, which regulate response to hypoxia^88^ and slow myosin expression^89,90^, respectively, are both crucial for embryonic development^91,92^, and were preferably upregulated in proximal muscles (**Supplementary Figure 4g**). Similarly, *METTL21C* was detected in and known to be restricted to *MYH7*-positive muscle fibers^93^, see **Supplementary Figure 4g**, while the expression of closely located *METTL21EP* has the opposite trend. This gene is considered non-functional in humans but encodes fast IIb myofiber-specific METTL21E in mice^94^ and is conserved across vertebrates^95^. Thus, we suggest its expression marks the remnant IIb muscle fibers, the glycolytic fibers associated with rapid contractile characteristics and high contraction forces, which were evolutionarily lost in humans^96^.

Considering fast- and slow-twitch muscles, we managed to distinguish them using a set of 42 marker genes from Abbassi-Daloii *et al.*^25^, which include slow *MYH7,* and fast *MYH1* and *MYH2*, see **Figure 2f**. We also recapitulated the observations of Moreno-Justicia *et al.*^76^: lncRNA *RP11-255P5.3* and *LINC01405* indeed serve as specific markers of fast and slow myofibers, respectively.

#### FANTOMUS uncovers expression patterns of genes related to muscular dystrophies

EOM are known to resist various muscular dystrophies^11^. This phenomenon has been studied at different levels, including unique eye muscle developmental origins^2^, innervation pattern^11^, stem cell niche^97,98^, and gene expression profile^12,99^, with the latter revealing EOM molecular phenotype as a combination of that of skeletal and cardiac muscle fibers. In turn, we focused on the genes whose mutations cause muscular dystrophies that spare EOM^100^. As such, both *CAPN3* and *DYSF*, associated with calpaino- and dysferlinopathies, respectively, were downregulated in EOM on the transcriptome and proteome levels (**Figure 2e**). Notably, in other muscles but not EOM, the abundance of dysferlin anticorrelated with CD55 (known to compensate for dysferlin deficiency^101^), see **Supplementary Figure 5d**. Dystrophin protein abundance was also lower in EOM, supporting their resistance to degeneration caused by DMD. However, dystrophin functional analogue utrophin (encoded by *UTRN*), thought to cause the observed resistance^102^, was upregulated on the RNA-level only (**Supplementary Figure 5d**). Recent studies found that both the *mdx:utrophin*+/− (*Dmd-/-:Utrn+/-*) and *mdx:utrophin−/−* (*Dmd-/-:Utrn-/-*) mice were morphologically spared in DMD^103^, suggesting there are other means of EOM resistance. One possible explanation could be the involvement of the vinculin-talin-integrin system, and we indeed found integrin beta-1 protein upregulation in EOM, which was shown to interact with a short dystrophin isoform in neuronal tissue^104^, while the double knockout β1KO*mdx* (*Itgb1-/-:Dmd-/-*) mice demonstrated more severe cardiac dysfunction compared to individual knockouts^105^ (**Supplementary Figure 5d**).

Besides EOM, other muscles also respond differently to various myopathies^106–108^. Thus, we curated a panel of 18 likely causative genes supported by MRI-based assessments of leg muscle fat replacement patterns (see **Methods** and **Supplementary Data 5**). Next, we classified muscles into three groups: typically spared in a particular myopathy, affected later, and affected earlier or severely. By comparing the expression level of each target gene among the specified groups, we observed a significant differential *FKRP* expression between these groups, i.e., between leg muscles differently affected by the FKRP-related dystroglycanopathy (ANOVA Holm-adjusted *P*-value = 0.005), **Figure 3a** and **Supplementary Data 5**. This aligns with the reduced selective pressure on FKRP according to gnomAD (v.4.1)^109^: *FKRP* MANE transcript has the lowest predicted loss-of-function (pLoF) z-score of -0.43 and is the only gene from the 9th loss-of-function observed/expected upper-bound fraction (LOEUF) decile among all myopathy-related genes included in our test. Moreover, the dystrophic phenotype caused by *FKRP* mutation can be restored by modulating its expression^110^. Together with the observations made for EOM, this case study highlights the value of FANTOMUS data in tracing the expression distribution of medically relevant causative genes across different muscles, paving the road to a better understanding of involved molecular mechanisms and predicting individual muscle susceptibility to particular myopathies.

**Figure 3.**
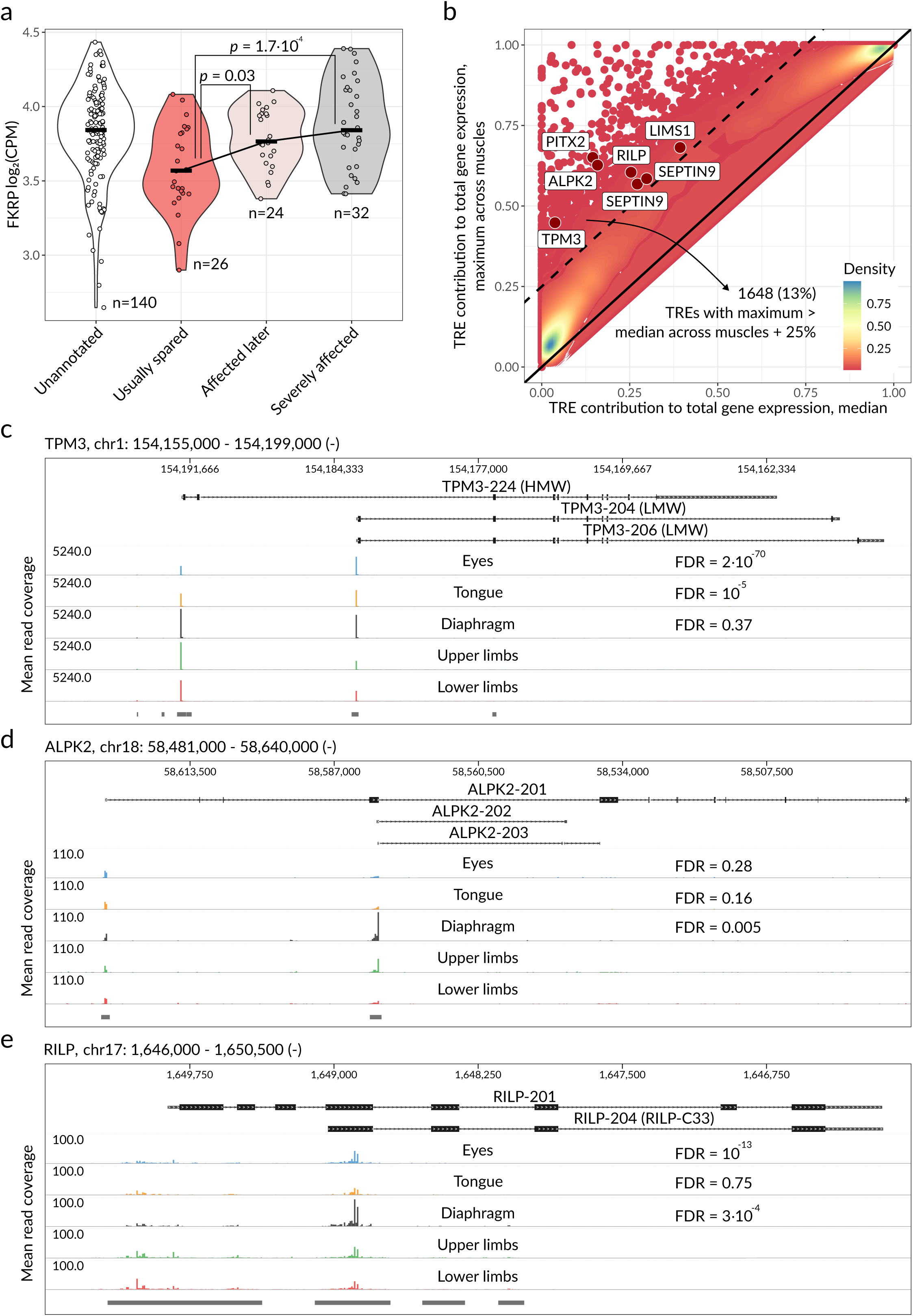
Differential *FKRP* expression and TRE usage distinguish muscles by susceptibility to dystroglycanopathy and body parts, respectively. **a**, *FKRP* expression (log_2_CPM) across muscles grouped by their susceptibility to dystroglycanopathy. Color: muscle group, bars: medians, n: sample sizes. *P*-values: Tukey’s honest significance test. **b**, Scatter and density plots for the maximum contribution of an individual TRE to the total gene expression (CPM) across muscles versus the median. Black solid line: y = x, dashed line: TREs with the maximal contribution 25% greater than the median. Color: density, examples discussed in the text are labeled. **c**, *TPM3* genomic view (top), mean CAGE-Seq profiles for EOM, tongue, diaphragm, upper and lower limbs (middle), and FANTOMUS TRE track for the corresponding strand (bottom). Color corresponds to the muscle group. FDR for EOM, tongue, and diaphragm was obtained by performing differential TRE usage analysis with edgeR. **d-e**, Differential TRE usage of *ALPK2* and *RILP* illustrated in the same way as in **(c)**. TRE: transcribed regulatory element, CPM: counts per million, FDR: false discovery rate, HMW: high molecular weight isoform, LMW: low molecular weight isoform.

#### Differentially expressed transcripts and protein isoforms arise from alternative promoter usage

In fact, CAGE-Seq measures not the total gene expression but the activity of individual TREs, enabling assessment of the alternative promoter usage. We focused on 5070 multi-TRE genes and a subset of 12928 TREs, each of which was either the major TRE (had the maximal activity across muscles) or had the activity comparable to the major TRE in at least one muscle; genes with a single unmatched major TRE were excluded (see **Methods**). We then characterized each of these TREs by the relative contribution to the total gene expression of the respective gene. For 1648 TREs, the maximal contribution to the gene expression across muscles was 25% greater than the median, i.e., reflecting the alternative promoter usage specific to particular muscles, and 1065 TREs corresponded to annotated MANE or GENCODE promoters (**Figure 3b** and **Supplementary Data 6**).

Next, we compared extraocular muscles, tongue, and diaphragm to all other samples and identified 4800, 534, and 896 genes, respectively, with significant differential TRE usage (edgeR diffSpliceDGE FDR < 0.05, see **Methods** and **Supplementary Data 7**). To illustrate the functional consequences, we selected 6 genes encoding transcription factors, protein kinases, and proteins involved in cytoskeleton organization (**Figures 3c-e** and **Supplementary Figure 6c**).

The first illustrative example is the tropomyosin gene *TPM3* encoding the striated high (HMW) and the cytoskeletal low molecular weight (LMW) isoforms. The former is known to tune the contraction of slow myofibers and, according to the FANTOMUS data, was downregulated in EOM, which are frequently spared in TPM3-related myopathy^111^, **Figure 3c**. Given that TPM3 was the most abundant protein in MS, we confirmed the observed tissue-specific TRE usage by distinguishing the intensities of its HMW and LMW isoforms, see **Supplementary Figure 6d**. Second, we focused on PITX2, the transcription factor involved in vertebrate embryonic development, and particularly important for heart formation^112–114^ and EOM specification^115–117^. *PITX2* mutations cause Axenfeld-Rieger syndrome, characterized by ocular, dental, and umbilical abnormalities^118^. Short *PITX2* isoform *PITX2C,* known to be involved in left-right body asymmetry^119^ and muscle regeneration^120–123^, was ubiquitously expressed across muscles. In turn, longer isoforms *PITX2A* and *PITX2B* demonstrated EOM-specific upregulation^124^ (**Supplementary Figure 6c,d**). The third example is *ALPK2*, which encodes the cardiac-specific atypical protein kinase previously found to promote cardiogenesis^125^ and prevent cardiac diastolic dysfunction, possibly through tropomyosin 1 phosphorylation^126^. We found the major *ALPK2* isoform to be ubiquitously expressed across muscles, while TRE, located near the fourth exon, was specifically upregulated in diaphragm and its neighboring muscles, see **Figure 3d** and **Supplementary Figure 6b**. Notably, *circALPK2* RNA generated by the fourth exon back-splicing was identified as a negative regulator of cell proliferation in human embryonic stem cell-derived cardiomyocytes via *circALPK2/miR-9/GSK3B* axis^127^.

Other examples of alternative TRE usage included *SEPTIN9, RILP,* and *LIMS1*, all encoding the cytoskeleton proteins. The former produces long isoform *SEPTIN9_i1,* which includes a microtubule-associated protein (MAP)-like motif^128^ and preferably localizes to the nuclear envelope, reducing its deformability and promoting juxtanuclear invadopodia^129^. In turn, the shorter isoform *SEPTIN9_i2* only interacts with actin filaments, demonstrates the opposite intracellular localization, and inhibits cell migration^130^. We observed eye-specific depletion of *SEPTIN9_i1* and the increased expression of shorter isoforms, including *SEPTIN9_i2,* in EOM and diaphragm, **Supplementary Figure 6c,d**. Given the critical role of *SEPTIN9* in regulating myoblast differentiation during the initial commitment phase^131^, the identified preferences in *SEPTIN9* isoform usage across muscles likely contribute to the difference in the overall regeneration capacity of skeletal muscles. *RILP* encodes the protein that affects vacuolar ATPase activity, interacts with Rab proteins, and recruits dynein-dynactin motor complexes, thereby regulating late lysosomal and endocytic transport, phagosomes fusion with lysosomes, and autophagy^132–135^. The truncated RILP protein form, RILP-C33, lacking the N-terminal half, was shown to inhibit epidermal growth factor and low-density lipoprotein degradation, cause dispersion of lysosomes, and prevent the fusion of phagosomes with late endosomes and lysosomes^136,137^. We found the full-length *RILP* isoform downregulated in eyes, tongue, and diaphragm, while a shorter isoform was overexpressed in the latter, **Figure 3e**. Finally, *LIMS1* (or *PINCH1*) is a gene encoding a protein involved in F-actin bundling and focal adhesion as a part of ILK-PINCH-PARVIN (IPP) complex^138^. We observed *LIMS1* to be downregulated in non-somitic muscles, including EOM, and tongue, while its short isoforms encoding the protein with the altered N-terminal sequence were specifically upregulated in diaphragm, see **Supplementary Figure 6c**. *LIMS1* was demonstrated to affect chondrogenesis^139^, mitochondrial fragmentation^140^, cell proliferation, and myoblast differentiation, with the double ablation of *PINCH1* and *PINCH2* leading to early postnatal lethality with reduced size of skeletal muscles and detachment of diaphragm from the body wall^141^.

### Motif activity response analysis reveals distinct gene regulatory programs across muscles

The transcriptomic differences between cell types are driven to a major extent by the cell type-specific activity of transcription factors. To identify the key regulators that define muscle molecular heterogeneity, we performed the motif activity response analysis, MARA, with MARADONER and 478 non-redundant motifs (a subset of HOCOMOCO v14 collection) recognized by 785 transcription factors, see **Methods**. As a result, 353 motifs showed non-uniform activity across muscles (ANOVA FDR < 0.05), clearly separating skeletal muscle groups on the PCA plot, see **Supplementary Figure 7a** and **Supplementary Data 8**. We then focused on 16 distinct groups formed by 65 motifs contributing at least 1% to the variance explained by the first 5 principal components, see **Figure 4a**.

**Figure 4.**
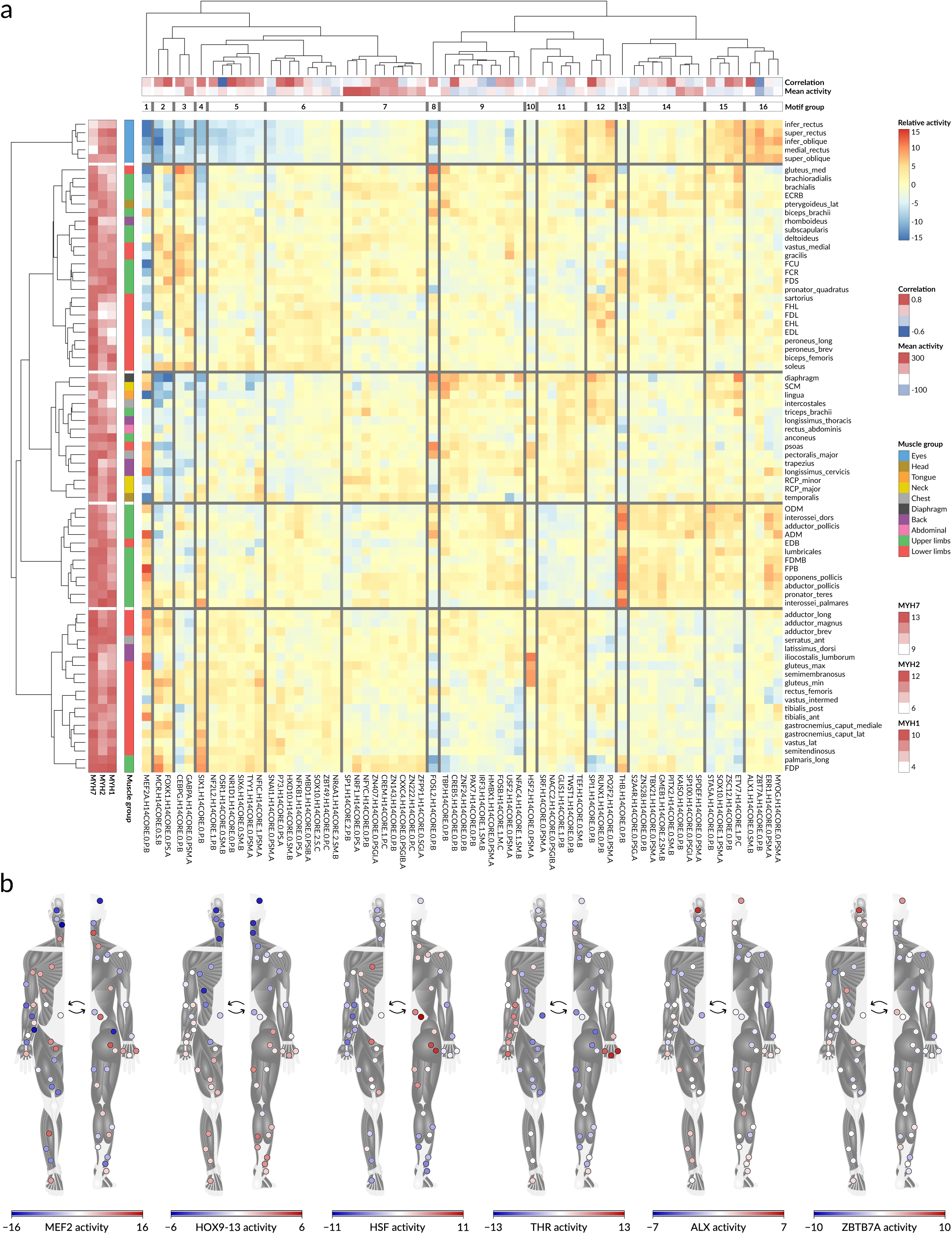
Transcription factor binding motifs with non-uniform activity across muscles. **a**, Motif activity heatmap across 75 skeletal muscles, 65 motifs contributing at least 1% to the variance explained by the first 5 principal components. Left: expression of slow- and fast-twitch MYH genes, log_2_CPM, and muscle group. Top: Pearson correlation coefficient between motif cluster activity and the total expression of the corresponding transcription factors, mean motif activity, and 16 distinct motif groups. Muscles and motifs were hierarchically clustered using Ward’s method with Euclidean distances. **b**, Left-to-right: MEF2, HOX9-13, HSF, THR, ALX, and ZBTB7A activities in different human skeletal muscles. Color: motif activity estimated by MARADONER. CPM: counts per million.

First of all, unique motif activity profiles were demonstrated by groups 1 and 4, which included MEF2 and SIX proteins. The former are preferably active in slow oxidative skeletal muscles^2^, but are also required for satellite cell differentiation and muscle regeneration^142^. The MEF2 group demonstrated non-trivial activity across muscles (**Figure 4b**), resembling the expression pattern of its direct target *CSRP3*^143^, **Supplementary Figure 7b**. *CSPR3* encodes a positive regulator of myogenesis and autophagy^144,145^, and its expression depends on MEF2 (Bayesian belief *p* = 0.963 for *m. flexor pollicis brevis*). In turn, the SIX proteins are the inductors of the fast-twitch phenotype^146,147^ and demonstrated upregulated activity in the corresponding muscles, see **Supplementary Figure 7c**. The motif group 5 included an alternative SIX motif, which demonstrated a similar activity pattern, while group 6 included the HOX9-13 motif cluster, which was preferably active in distal arms and legs, in agreement with our findings from TRE activity, see **Figure 4b**.

Besides MEF2 and SIX, other groups of singleton motifs included JUN/FOS proteins (group 8), heat shock factors (HSFs, group 10), and thyroid hormone receptors (THRs, group 13). The former, together with the larger motif groups 9, 11 and 12, were more active in slow-twitch muscles and included transcription factors involved in hematopoiesis and inflammation (SPI1, POU2, IRF3, RUNX), modulator of endothelium angiogenic potential ZNF24^148^, progenitor cell markers TWIST1^149^ and PAX7^150^, and pro-adipogenic transcription factor GLIS1^151^, possibly reflecting upregulated satellite cells proliferation, increased regenerative potential and adipogenesis of oxidative muscles^152–154^, see **Supplementary Figure 7c**. However, the expression levels of *TWIST1* and *PAX7 per se* were higher in EOM, which were also true for *FOXC1* and neuromuscular junction-associated *HSD11B1*, but not for *MYOG* and *MYOD1*, confirming previous findings regarding the specific status of the extraocular progenitor cells^98,155,156^, **Supplementary Figure 7d**.

On top of that, HSF proteins demonstrated an increased motif activity in the hip and back specifically, **Figure 4b**, that correlates with the *HSPA6* expression pattern (Bayesian belief *p* = 0.976 for *m. iliocostalis lumborum*), **Supplementary Figure 7b**. These transcription factors are known to regulate the expression of heat shock protein genes in response to muscle damage and were found to be upregulated during aging^157,158^. Together with the elevated susceptibility of the trunk and thigh muscles to age-related atrophy^159^, these differences likely reflect the consequences of a sedentary lifestyle.

Thyroid hormone receptors had remarkable specificity towards distal arm muscles, which were previously found to express IIb marker *METTL21EP*, see **Figure 4b** and **Supplementary Figure 4g**. The expression pattern of hairless homolog *HR*, a well-known direct target of THRs^160,161^, resembles the observed differential activity of this motif cluster (Bayesian belief *p* = 0.989 for *m. opponens pollicis*), see **Supplementary Figure 7b**. Moreover, *MYLK4* promoter likely contains occurrences of the same motif, and its expression in somitic muscles also correlated with THRs activity (Bayesian belief *p* = 0.926 for *m. opponens pollicis*, motif *p* = 1.1*10^-6^), **Supplementary Figure 7b**. This gene encodes a myosin light chain kinase, which positively regulates muscle strength and stiffness^162^ and is overexpressed after exercise^163^.

Finally, group 16 contained transcription factors with increased activity in EOM. Of them, ALX1/ALX4 and ZBTB7A targets were upregulated in the eye muscles exclusively, see **Figure 4b**. ALX was shown to regulate periocular mesenchyme development^164^, ocular vascularization^165^, and muscle stem cell functioning^98^. ZBTB7A is a transcriptional repressor associated with macrocephaly and intellectual disability^166^, which was found to induce epithelial–mesenchymal transition of lens epithelial cells by activating Wnt/β-catenin signaling^167^ and to promote a switch from fetal to adult hemoglobin^168^. We identified *MYH15* and *ATP1A3* promoters as ALX and ZBTB7A targets, respectively (Bayesian belief *p* = 0.804 and 0.947 for *m. obliquus inferior*, motif *p* = 1.1*10^-4^ and 3.8*10^-6^), see **Supplementary Figure 7b**. *MYH15* is a myosin heavy chain gene expressed exclusively in the extraocular muscles^169^, while *ATP1A3* was found previously to be expressed in neurons, heart and EOM^170^, is enriched in neuron-derived extracellular vesicles^171^ and mutations in this gene are associated with Rapid-onset Dystonia-Parkinsonism (RDP), Alternating Hemiplegia of Childhood (AHC), Cerebellar ataxia, Areflexia, Pes cavus, Optic atrophy, and Sensorineural hearing loss (CAPOS) syndrome, and developmental and epileptic encephalopathy^172–174^.

### Allele-specific analysis reveals regulatory SNPs affecting muscle TRE activity

In addition to the gene- and TRE-level expression, CAGE-Seq allows for capturing the effects of regulatory single-nucleotide variants^47,175^ using allele-specific analysis. We called 16205 sufficiently covered single-nucleotide polymorphisms (SNPs) directly from the CAGE-Seq reads and estimated the allelic imbalance using MIXALIME^47^, see **Methods**. This resulted in 6653 allele-specific variants (ASVs) passing FDR < 0.05 in at least one muscle, **Supplementary Data 9**. On average, 60% of the identified ASVs were annotated as skeletal muscle expression quantitative trait loci (eQTLs) in GTEx (v.10)^54^; 54% coincides with allele-specific transcription factor binding sites (ASB) of ADASTRA (v.6.1)^176^; 10% ASVs coincided with muscle chromatin accessibility-altering variants according to UDACHA (v.1.0.3)^47^; 7% overlapped the skeletal muscle expression eQTLs reported in Wilson E.P. *et al.*^177^, see **Supplementary Figure 8a**.

To identify the functional relevance of the detected ASVs, we selected 58 skeletal muscle-related GWAS summary statistics available at NHGRI-EBI GWAS Catalog^178^ and Pan UK Biobank (https://pan.ukbb.broadinstitute.org), including appendicular lean mass, thigh muscle volume, hand grip strength, impedance, total weight, and others, see **Supplementary Data 9**. From those, we excluded the phenotypes with low total SNP-heritability and kept only one trait of each pair with high genetic correlation, as recommended by Finucane *et al.*^179^. For the resulting set of 32 analyzed phenotypes, we performed the partition heritability analysis (**Supplementary Figure 8b**), which revealed the significant enrichment of ASVs for the total thigh muscle volume z-score from van der Meer *et al.*^33^, see **Figure 5a**, suggesting the polygenic contribution of TRE-located ASVs to the heritability of the skeletal muscle volume.

**Figure 5.**
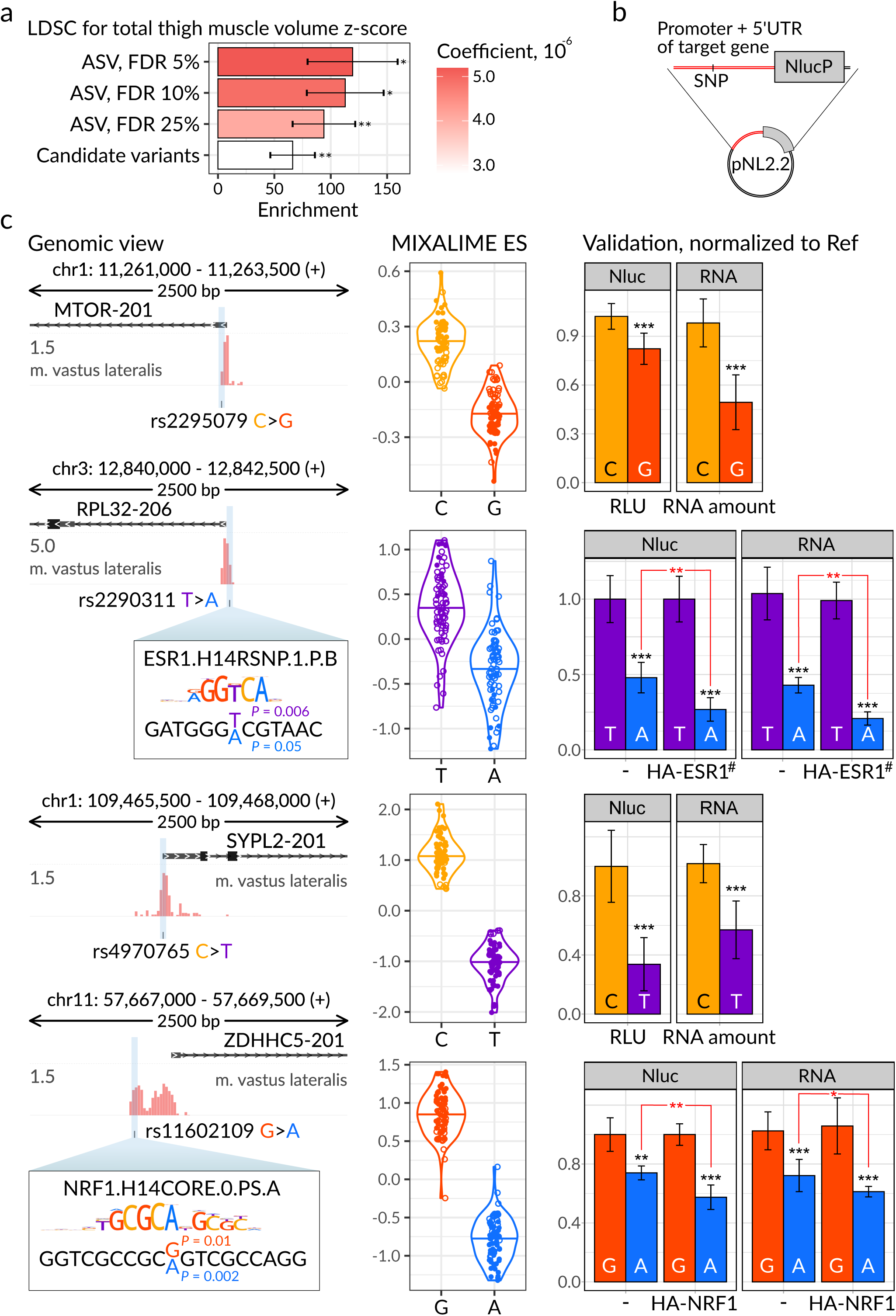
Regulatory SNPs contribute to skeletal muscle gene expression. **a**, The results of stratified LD score regression for total thigh muscle volume z-score from van der Meer *et al.*^33^ applied to all candidate sequence variants or only those passing certain allelic imbalance significance thresholds. Color: regression coefficient, error bars: standard errors, the enrichment *P*-values from sLDSC were FDR-adjusted for the number of tested traits. *FDR < 0.05, **FDR < 0.01. **b**, The scheme of the reporter vector. A native promoter and 5’ UTR region were cloned into the promoter-less pNL2.2 vector, encoding the NanoLuc luciferase fused with the PEST domain. **c**, Allele-specific variants selected for validation. Left: genomic view of *m. vastus lateralis* CAGE-Seq profile and the sequence variants; Y-axis: log_2_ mean number of 5’ ends of reads. For rs2290311 and rs11602109, the transcription factor binding motifs from HOCOMOCO v14 are shown above the binding site sequences; the motif *P*-values are labeled. Middle: sample-level allelic imbalance (log_2_) between the reference and alternative alleles, samples with FDR < 0.05 are shaded, bars represent the medians. Right: The results of luciferase reporter analysis with the luciferase activity (left) and reporter mRNA level (right) measured for the reference and alternative alleles, normalized to the reference allele. For rs2290311 and rs11602109, the HEK293T cells were transfected with the mixture of pNL2.2 and the relevant TF-overexpressing pcDNA3.1-HA vector to estimate the effect of ESR1 and NRF1 overexpression, respectively. Values represent the means of at least three independent replicates; error bars: standard deviation. *P*-value: two-tailed Student’s t-test. \**P*-value < 0.05, \*\**P*-value < 0.01, \*\*\**P*-value < 0.001. LDSC: linkage disequilibrium score regression, ASV: allele-specific variant, SNP: single-nucleotide polymorphism, FDR: false discovery rate, UTR: untranslated region, ES: effect size, Ref: reference allele, RLU: relative luciferase units, ESR1^#^: constitutively active ESR1 (Y537S).

Finally, we selected a small set of ASVs for validation in a luciferase reporter assay. To this end, we chose 4 variants demonstrating consistent allelic imbalance across different muscles, located within promoter TREs, and annotated as skeletal muscle eQTLs in GTEx, ASBs in ADASTRA, or associated with muscle-related phenotypes, see **Supplementary Data 9**. We cloned native promoters and 5’ UTRs into a promoter-less reporter vector, pNL2.2 (**Figure 5b**), for both allelic variants, and assessed the impact of the SNVs on reporter mRNA expression by measuring luciferase activity or performing qPCR in HEK293T cells. To confirm the involvement of ESR1 and NRF1 TFs in allelic imbalance at rs2290311 and rs11602109 (according to ADASTRA and motif analysis), we also repeated the assays in cells overexpressing the respective TFs.

The results shown in **Figure 5c** included rs2295079 and rs2290311 SNPs located in promoters of *MTOR* and *RPL32*, respectively, which are involved in muscle response to exercise and growth^180–182^. Both variants are annotated as eQTLs in various tissues, including skeletal muscles; rs2295079 is associated with the body fat-free mass and weight, and rs2290311 was found to significantly affect estrogen receptor binding (**Figure 5c**, middle panel). Considering the estrogen’s role in promoting skeletal muscle stem cell differentiation^183^ and preventing sarcopenia^184^, the observed ESR1-dependent allelic imbalance likely contributes to the maintenance of the total skeletal muscle mass.

The other two validated variants, rs4970765 and rs11602109, are located in the promoters of *SYPL2* and *ZDHHC5*, which are involved in muscle differentiation through t-tubule formation^185,186^ and lamin A palmitoylation^187^, respectively. These variants are associated with the body impedance and fat-free mass, and are annotated as skeletal muscle eQTLs. Moreover, rs11602109 was found to significantly affect the binding of nuclear respiratory factor-1, which is a known regulator of mitochondrial biogenesis^188^ and glucose transport^189^ (**Figure 5c**, bottom panel). Dysregulation of *ZDHHC5* and *SYPL2* is associated with heart failure^190^, sarcopenia^191^, and even muscular dystrophy^192^; thus, rs4970765 and rs11602109 likely directly affect the molecular pathways involved in muscle functioning.

### Limitations of the study

Collecting a systematic and diverse set of biopsies from skeletal muscles of a single healthy individual is unrealistic, and FANTOMUS data were obtained from autopsy samples of only four post-mortem donors. Thus, the resulting CAGE-Seq profiles might be incomplete or biased, e.g., due to individual physiological state of particular donors (such as reduced muscle activity) or systematic RNA degradation, although the RNA integrity was acceptable (**Supplementary Data 1**). To further clarify the FANTOMUS reliability, we quantitatively compared FANTOMUS *m. soleus* TREs to the CAGE-Seq data from Bokov *et al.*^193^ obtained from the biopsy samples of the same muscle, which resulted in PCC of 0.91 (**Supplementary Figure 3d**).

While FANTOMUS was successful in capturing the activity of promoters (**Figure 1d**, **Supplementary Figure 1e**), it had limited versatility in identifying bidirectional enhancers. In total, we annotated only 881 actively transcribed enhancers compared to 7554 detected as active in at least one of 19 FANTOM5^49^ skeletal muscle samples. However, if we exclude the primary cells derived from embryos and fetal muscles, FANTOM5 muscle samples cover only 6 adult skeletal muscle tissues (including one undefined, 4 EOM, and one *m. soleus* sample) with only 1471 actively transcribed enhancers; thus, the rest likely represent the unique enhancer repertoire of the fetal tissues and myotubes differentiated *in vitro*^194^. Requiring the FANTOM5 enhancers to be reproducibly detected between samples, i.e., in both adult skeletal muscles or in all four EOM, the list shrinks to 125 TREs only, of which only 93 (90 autosomal) pass the CPM > 0.5 threshold in all EOM or adult skeletal muscle samples. Of those 93, 70 are recovered by FANTOMUS, 60 by uni-, and 12 by bidirectional TREs; the latter by design do not overlap any known gene TSS from MANE and GENCODE, and demonstrate the reproducible bidirectional activity across skeletal muscles. All in all, the FANTOMUS enhancer catalog is most likely incomplete, but is comparable in scale to FANTOM5 adult muscle data, likely reflecting the limitation of CAGE-based detection of transcribed enhancers in adult skeletal muscle samples.

Lastly, FANTOMUS was built upon bulk CAGE-Seq and MS data and therefore can not distinguish the molecular profiles of individual fibers within the analyzed skeletal muscles. While numerous studies focused on single-cell and single-nucleus transcriptomics and proteomics have already been published for a small subset of skeletal muscles^20,21,24,76^, studying both intra- and inter-muscular molecular heterogeneity remains an untackled challenge.

### Concluding remarks

FANTOMUS atlas highlights significant heterogeneity of the skeletal muscle molecular profiles on both transcriptome and proteome levels: about 80% of genes and proteins are differentially expressed across muscles, and more than 20% have a strong preference towards specific muscles. The correlation of the promoterome and proteome is limited but stable across diverse muscles (SCC around 0.22), with EOM being an outlier with SCC of 0.34, likely reflecting a relaxed post-transcriptional expression control.

Both transcriptome and proteome data highlighted EOM, tongue, and diaphragm as muscles with the most outlying molecular profiles. However, other muscles also have distinct expression profiles and can be distinguished, e.g., based on HOX gene expression, fast and slow-twitch fiber markers, or *FKRP* expression, with the latter separating tissues by their involvement in the FKRP-related dystroglycanopathies.

Taking advantage of the CAGE-Seq data, we managed to dissect distinct skeletal muscle regulatory programs beyond the total gene expression estimates. FANTOMUS revealed a widespread tissue-specific alternative TRE usage, including the key EOM development regulator PITX2, cardiac-specific protein kinase ALPK2, sarcomere component TPM3, and other cytoskeleton proteins. MS data confirmed EOM-specific downregulation of TPM3 HMW isoform, which might explain the resistance of these muscles to TPM3-related myopathy.

In addition to the differential TRE activity *per se*, we estimated the activities of the transcriptional regulators that drive the observed expression differences. MARA identified more than three hundred transcription factor motifs with non-uniform activity across muscles. We identified multiple TF motifs preferably active in slow, fast, and extraocular muscles. Further, besides muscle heterogeneity, we studied the differences in TRE activity caused by individual genomic variation. For this, we called more than 16 thousand SNVs and performed the allele-specific analysis, which resulted in 6,6 thousand ASVs. The identified variants demonstrated the enrichment for the loci associated with total thigh muscle volume z-score and were selectively validated in a luciferase reporter assay.

Finally, we developed a dedicated web portal to the FANTOMUS data accessible at https://fantomus.autosome.org, which provides interactive access to the expression profiles of 18 thousand genes, abundance estimates of 1,8 thousand protein groups, and activity of 37 thousand TREs and 478 transcription factor motifs across 75 distinct human skeletal muscles. We envision FANTOMUS as a multimodal hub of human skeletal muscle molecular phenotypes and believe that these data will facilitate further insights into the regulatory logic distinguishing human skeletal muscles in normal and pathological conditions, including susceptibility to myopathies.

## Methods

### Ethical Statement

The study was approved by the Institute of Protein Research’s ethical committee as complying with relevant ethical regulations and was determined as non-human subject research. De-identified muscle tissue samples were procured by "Natsionalny BioService" LLC (St.Petersburg, Russia) in accordance with the Russian Government Resolution No. 750 of July 21, 2012, "On Approval of the Rules for the Transfer of the Unclaimed Body, Organs, and Tissues of a Deceased Person for Use in Medical, Scientific, and Educational Purposes, as well as the Use of the Unclaimed Body, Organs, and Tissues of a Deceased Person for these Purposes" (http://pravo.gov.ru/proxy/ips/?docbody=&nd=102158324&rdk=&backlink=1) in accordance with Russian legislation.

### Skeletal muscle sample preparation and CAGE-Seq

#### Sample preparation

In the presented study, we used 228 autopsy tissue samples from 4 individuals (3 males and 1 female, deceased due to hemorrhagic stroke or coronary heart disease, 56-67 years old), including 3 non-muscle *septum intermusculare brachii laterale* (used only for variant calling), see the "Allele-specific analysis" section. The samples were fixed in RNA-later solution (Invitrogen), stored at +4 °C overnight, and then stored at -80 °C until RNA isolation. Total RNA was extracted from muscle samples using the RNeasy Fibrous Tissue Kit (Qiagen, Germany) according to the manufacturer’s protocol with minor modifications. 19-30 mg of tissue was homogenized with a mortar and pestle using liquid nitrogen. After that, the lysis buffer was added, and the lysate was collected into clean tubes. The samples were then incubated with proteinase K solution for 10 minutes at 55°C and centrifuged in an Eppendorf mini spin centrifuge at maximum speed. To precipitate long and short RNAs, one volume of 96% ethanol was added to the supernatant, and all was applied to a column. The column was washed twice with RPE wash buffer, dried, and eluted with nuclease-free water.

#### Library preparation and sequencing

Libraries were prepared utilizing the standard nAnT-iCAGE (non-Amplified non-Tagging Illumina Cap Analysis of Gene Expression) protocol described in Murata *et al.*^195^. 5 μg of total RNA was used as a template for synthesis of the first cDNA strand (nAnT-iCAGE Library Preparation kit DNA form, Yokohama, Japan, and SuperScript III Reverse Transcriptase, Invitrogen, Waltham, MA, USA). Next, RNA/DNA heteroduplexes were biotinylated at the 5’-ends (nAnT-iCAGE Library Preparation kit, DNAform, Japan), allowing selection of 5’-cap containing RNAs using streptavidin beads (Dynabeads M-270 Streptavi din, ThermoFisher Scientific, USA). Then, cDNA was treated with RNase I and H (nAnT-iCAGE Library Preparation kit, DNA form, Japan) and purified using RNACleanUP (Beckman Coulter, Brea, CA, USA). Next, linkers were ligated to the 5’- and 3’-ends (nAnT-iCAGE Library Preparation kit, DNAform, Japan) of the cap-trapped cDNA. In the final stage, a second cDNA strand was synthesized from these short CAGE tags (nAnT-iCAGE Library Preparation kit, DNAform, Japan). The concentration of the resulting libraries was determined by the PicoGreen Assay in a GloMax® Multi Detection System (Promega, Madison, WI, USA), and the libraries were validated using real-time PCR (KAPA Library Quantification Kits Illumina, KAPA Biosystems, Wilmington, MA, South Africa). The sequencing was performed on either Hiseq2500, Nextseq 550, or Novaseq6000 platform (Illumina, San Diego, CA, USA)

### CAGE-Seq data processing

#### Preprocessing and quality check

The resulting fastq files from 225 muscle samples contained from 10 to 100 million reads and were aligned to the hg38 genome assembly using HISAT2 (v.2.2.1)^196^ with GENCODE comprehensive annotation (v.44)^197^ and a very-sensitive preset. For allele-specific analysis, the fastq files were additionally pre-processed using cutadapt (v.4.1) `-u 3 -q 20 -m 20 -a AGATCGGAAGAG -a CACACGTCTGAACTCCAGTCAC` to remove short sequences and trim low-quality ends, Illumina adapters, and the first 3 bases from the 5’ end, which often contain unencoded guanines^198,199^ (see below). Next, we filtered out multimapped reads with samtools (v.1.13) `view -q 60` and performed gene counting with featureCounts (v.2.0.6) `-s 1` for sample quality check. This included PCA based on the uncorrected log_2_CPM values using R (v.4.3.1) `prcomp` from stats and factoextra (v.1.0.7), as well as bulk data deconvolution to predict the cell type composition. The deconvolution was performed using SKM_human_pp_cells2nuclei_2023-06-22.h5ad from Kedlian *et al.*^21^, zellkonverter (v.1.10.1), Biobase (v.2.60.0), and SingleCellExperiment (v.1.22.0) R packages for data manipulation and `music_prop` from MuSiC (v.1.0.0)^200^ to calculate cell type fractions, see **Supplementary Data 1**. Together with the total allelic imbalance, see the "Allele-specific analysis" section, these steps allowed us to identify and remove 3 potentially mixed-up or contaminated samples, namely *lingua* from the individual 5708, which was mislocated on the PCA and demonstrated low myofiber content, *m. abductor digiti minimi* from 5708 and *m. pterygoideus lateralis* from 5489, which showed the total reference bias, defined as the fraction of variants with the reference > alternative allelic read coverage, of 0.55 and 0.56, respectively, compared to ∼0.43 for the rest of the samples, see **Supplementary Figure 1b-d**. This resulted in 222 muscle samples being processed further with samtools, awk (v.5.1.0), bedtools (v.2.30.0) `genomecov -bg -5`, and bedGraphToBigWig (v.2.10) to generate bedGraph and bigWig tracks of the 5’-ends of all reads (allN), or starting from one to three prepended "soft-clipped" guanines (softG). These guanines usually remain unaligned to the genome sequence, as the corresponding cytosines were externally added to the cDNA by the reverse transcriptase reading through the cap structure^198,199^. Thus, their presence is considered a "cap signature" marking high-confidence CAGE-Seq reads, see also the next section.

#### Peak calling

The peak calling was performed with CAGEfightR (v.1.22.0)^201^. Normally, the high-confidence CAGE-Seq reads should be marked by softflipped 5’ Gs from reverse transcription. However, a single G or a stretch of Gs can occur upstream in the genome sequence directly adjacent to a TSS. Consequently, in this case, the sequenced guanines incorporated by the reverse transcription will yield a perfect (unclipped) alignment. To properly account for the TSS-adjacent nucleotides, we used a hierarchical filtering and annotation procedure for prioritizing high-confidence TREs, see **Supplementary Figure 2a**:

1. Filtering genomic positions, CAGE transcription start sites (CTSS), covered by read 5’-ends (CAGE tags) in 3 or more samples, leaving 15177664 allN and 4495316 softG CTSS;
2. Сalculating the difference between allN and softG profiles to obtain non-clipped position coverage (noclip);
3. Filtering out softG for 5’-G[N]-3’ (leaving 4495312 softG CTSS) and noclip for 5’-N[G]-3’ (leaving 6732031 noclip CTSS) to prevent inflating the noclip TRE coverage by non-encoded 5’ G accidentally aligned to the reference genome;
4. Identifying bidirectional CTSS clusters, i.e., bidirectional TREs, with `clusterBidirectionally`, `calcBidirectionality`, and `quantifyClusters` for 17470 TREs demonstrating transcription in both directions in at least one sample;
5. Annotating bidirectional TREs with MANE (v.1.1, first round) and GENCODE (v.44, second round for TREs left unannotated) using `assignGeneID` and `assignTxType` for each strand separately, followed by filtering out TREs annotated as promoters either on forward or reverse strands (14969 bidirectional TREs remain);
6. Identifying unidirectional TREs with `clusterUnidirectionally` and `quantifyClusters` using CTSS, which remained after excluding bidirectional TREs (2593090 unidirectional TREs);
7. Combining 2608059 bi- and unidirectional TREs and annotating with MANE (v.1.1, first round) and GENCODE (v.44, second round for TREs left unannotated or annotated by MANE as non-promoter regions of the same gene);
8. Leaving only 1360769 verifiable TREs, i.e., those containing softG or noclip CTSS from (3);
9. Reducing the TRE list to 37186 credible regions that pass either "soft-clipped CPM > 10" or "total CPM > 0.5 and soft-clipped CPM fraction ≥ 0.5" in 3 or more samples.

The full script is available at https://github.com/autosome-ru/fantomus-clustering.

A two-step annotation procedure was performed to prioritize high-confidence transcripts from MANE over others (e.g., miRNA genes). A threshold for the minimal soft-clipped CPM fraction was set to mostly preserve the promoter regions, see **Supplementary Figure 2b**. Batch-correction was performed with ComBat-ref^202^ (https://github.com/xiaoyu12/Combat-ref, accessed in March 2025) specifying different sequencing platforms and runs (sequencing batches) as batches and muscles as a biological covariate. The last step included filtering CAGE-Seq credible clusters potentially originating from the "exon painting"^203,204^, which was performed using bedtools `intersect -s -f 0.95 -r -v` to remove clusters intersecting CDS from MANE (v.1.1) and GENCODE (v.44) by 95%. This procedure resulted in the final set of 37001 credible FANTOMUS TREs. To obtain gene-level expression estimates, we summed up cluster counts of 32433 annotated unidirectional TREs into 18329 genes.

#### Visualization

Data visualization was performed in R using ggplot2 (v.3.5.0) and pheatmap (v.1.0.12) packages. To generate the genomic view of the CAGE-Seq profiles (**Figures 1c** and **3c-e**,

**Supplementary Figure 6c**), we used bedGraph files containing reads starting with soft-clipped G, and passed them to svist4get (v.1.3.1)^205^ along with the GENCODE (v.44) annotation.

### Mass spectrometry

#### Sample preparation

For each of 60 muscle samples selected for MS, see **Supplementary Data 1**, a piece of frozen tissue was transferred to a liquid nitrogen-cooled mortar and ground with a pestle to obtain fine tissue powder. Five to ten mg of powder was transferred to 135 μl of lysis buffer (5% sodium dodecyl sulfate, 25 mM dithiothreitol, 50 mM HEPES, pH 8.2) and vortexed. The lysate was transferred to an AFA microtube, sonicated (average power 20 W, 2 min × 2) using a ME220 sonicator (Covaris, USA), incubated at 55°C (15 min), and centrifuged (5 min, 30,000 g). The proteins (140 μg, fluorimeter Qubit 4, Thermo Fisher Scientific, USA) were diluted in lysis buffer (final volume 24 μl), then 1 μl of 500 mM iodoacetamide was added, and the mix was incubated in the dark (10 min). Each sample was treated according to the S-Trap micro column digestion protocol (ProtiFi, USA) using trypsin and Lys-C in 25 μl 50 mM HEPES, pH 8.2 (recombinant modified Trypsin-MS-RS [1:22], Molecta, Russia, and Lys-C VA1170 [1:112], Promega, USA, accordingly; 8 h at 37 °C). After elution, 10 μg of peptides (fluorimeter Qubit 4) were dried, resuspended in 200 mM HEPES, pH 8.2, and labeled with TMT 16-plex isobaric labels (Thermo Fisher Scientific) in 40% acetonitrile for 1.5 h. The reaction was stopped (0.3% hydroxylamine w/v, 15 min), and samples were pooled. The mixture of labeled peptides was separated using high pH reverse-phase LC fractionation (HPLC 1200, Agilent, USA). The peptides were concentrated on an analytical column (XBridge C18, particle size 5 μm, 4.6 x 250 mm, Waters, Ireland) in isocratic mode at a flow of 750 μl/min for 3 min in mobile phase A (15 mM ammonium acetate, pH 9,0). Twenty-four fractions were collected from 3 min to 50 min (collection time 2 min, volume 1500 μL) using a gradient elution mode with mobile phase B (15 mM ammonium acetate, pH 9.0, 80% acetonitrile, pH 9.0). Each fraction was concentrated, and then the fractions were combined to obtain 10 resulting fractions (1, 2+3+13, 4+5+14, 6+15, … 11+20, and 12+21+22+23+24). Each fraction was analyzed two times using an HPLC Ultimate 3000 RSLC nanosystem (Thermo Fisher Scientific, USA) and a Q Exactive HF-X Hybrid Quadrupole-Orbitrap mass spectrometer (Thermo Fisher Scientific, USA) by the nanoelectrospray ion source in the positive mode of ionization (Thermo Fisher Scientific). The gradient (90 min) was formed by the mobile phase A (0.1% formic acid) and B (80% acetonitrile, 0.1% formic acid) at a 0.4 μL/min flow. The ionizing voltage was 2.2 kV. MS spectra were acquired at a resolution of 60,000 in the 400–1400 m/z range; fragment ions were mass scanned at a resolution of 60,000 in the range from m/z 120 to the upper m/z value as assigned by the mass to charge state of the precursor ion. All tandem MS scans were performed on ions with a charge state from z = 2+ to z = 4+. Synchronous precursor selection facilitated the simultaneous isolation of up to 10 MS2 fragment ions. The maximum ion accumulation times were set to 50 ms for precursor ions and 30 ms for fragment ions. AGC targets were set to 10^6^ and 10^5^ for precursor ions and fragment ions, respectively.

#### Estimating protein abundances

Peptide and protein/protein groups identification and search were conducted using the MaxQuant platform (v.2.7.5.0; Max Planck Institute for Biochemistry) using default settings (FDR for peptides 1%, N-terminal acetylation and oxidation of methionine as variable modifications and carbamidomethylation of cysteine, as fixed modification) and the Isobaric match between runs and PSM-level weighted ratio normalization functions^206^ with reference channels 10 to 15 for TMT-set 1 and 1 to 15 for TMT-set 2, 3, and 4. To search for protein isoforms encoded by mRNA from an alternative promoter, a protein sequence FASTA file (all reviewed human Swiss-Prot entries) was expanded with the sequences of the investigated isoforms. Further analysis was performed using a Perseus platform (v.2.0.11; Max Planck Institute for Biochemistry). After filtration (potential contaminants, reverse peptides, peptides identified only by site), proteins identified by more than one peptide (unique+razor) and presented in 75% of the samples were selected for further analysis. Reporter ion intensities were log-transformed and normalized to the sample medians. As a final step, we added a constant c to each value to preserve the original scale, where c was estimated as mean unnormalized log-transformed intensity across all protein groups and samples.

Further MS data analysis included annotating proteins by gene ENSEMBL ids with `mapIds` from AnnotationDbi (v.1.62.2) and org.Hs.eg.db (v.3.17.0), followed by intersecting with the CAGE-Seq results for 1750 autosomal genes to exclude proteins encoded in the mitochondrial genome and avoid sex bias. Deming regression for the log_2_(iBAQ from MS) and log_2_(mean CPM from CAGE-Seq) values, calculated for the same sample set, was performed using `mcreg` with `method.reg = ’Deming’` from the mcr (v.1.3.3.1) package. GSEA for the residuals was performed using fgsea (v.1.26.0)^207^ R package and the gene sets from MSigDB (v.2024.1)^208–210^. Finally, we calculated gene-level z-scores for the log_2_Intensity (reporter ion intensity from MS) and log_2_CPM (CAGE-Seq) values to calculate Pearson correlation coefficients for the individual muscle samples.

### Differential expression and TRE usage analysis

#### Assessing differential expression across muscles

To identify TREs, genes, and proteins that are differentially expressed or abundant across muscles, we used edgeR (v.3.42.4) for CAGE-Seq cluster- and gene-level read counts and limma (v.3.56.2) for protein-level log_2_Intensity estimates. For CAGE-Seq, we performed TMM-normalization with `calcNormFactors`, followed by `estimateDisp`, `glmQLFit`, and `glmQLFTest` as a part of edgeR’s quasi-likelihood pipeline, using "∼ individual + muscle" design to block by individuals. For protein abundance, we used `lmFit` with "∼ muscle" design, blocking by individuals, and the intra-block correlation obtained from `duplicateCorrelation`, followed by moderated statistics calculation using `eBayes`. To obtain the results of the pairwise comparisons between the particular skeletal muscle and all other samples, we used `topTags` and `topTable` for edgeR and limma, respectively, while ANOVA-like analysis was performed by setting the `coef` in `glmQLFTest` (edgeR) or `topTable` (limma) to a vector of coefficients corresponding to different muscles. The alternative promoter usage analysis of 32433 unidirectional TREs of 18329 genes was performed in edgeR in the same way, with the last two steps relying on `diffSpliceDGE` and `topSpliceDGE`. The significance of the TPM3 isoforms’ differential abundance was calculated separately with aov(log_2_Intensity ∼ individual + muscle) from stats (v.4.5.2). Causative gene differential expression across muscle groups with different susceptibility to the corresponding myopathy was estimated with aov(log_2_CPM ∼ individual + muscle group) followed by Holm *P*-value adjustment for the number of tested genes.

Skeletal muscles with the most uniquely expressed TREs were identified by GSEA-like analysis (with the fgsea R package) as those significantly overrepresented at the top after ordering TREs by the difference between the maximum and the median log_2_CPM across muscles, and assigning each TRE to a muscle where the maximum is achieved.

#### Estimating relative TRE contribution to gene expression

To visualize the relative contribution of individual unidirectional TREs to the total gene expression, we calculated the median CPM for each muscle, separately for each TRE. Next, only 1490887 TRE-tissue pairs for 21235 TREs of 7237 genes with at least 0.5 median CPM were left for a reliable TRE contribution estimation, which was calculated in each muscle as a fraction of TRE CPM divided by the gene-level sum. Technically, differential expression of a single highly-expressed promoter of a major transcript isoform could yield multiple detectable "promoter switching" side-effects, due to the increased relative contribution of minor isoforms to the total expression upon the downregulation of the major isoform. In such cases, the functional consequences are likely limited to the changes of the major isoform, as the effect of the differential expression can be of a magnitude higher than the alterations in the relative promoter activity and isoform balance. Thus, in our analysis, the final step included filtering out TREs for which the maximal expression across muscles, log_2_CPM, deviated from the maximal expression of any other TRE of the same gene by more than 5 in an absolute scale, see **Supplementary Data 6**, i.e., excluding the alternative isoforms with a negligible relative expression or single major isoforms with no comparable alternatives.

### Motif activity response analysis

For motif activity response analysis, we used MARADONER^211^ (MARA-done-right, v.0.25.3, https://github.com/autosome-ru/MARADONER), provided with an annotation of biological replicates for each muscle, log_2_CPM expression estimates for 36120 unidirectional TREs in 222 samples, and sum-occupancy scores^212^ of HOCOMOCO v14^213^ motifs calculated using SPRY-SARUS^214^ (v.2.0.2, https://github.com/autosome-ru/sarus) for the same TREs trimmed to 250 nucleotides upstream and 10 nucleotides downstream of the summit. Basically, MARADONER shares the original logic of MARA^215^ and isMARA^216^ with an assumption that the promoter activity in each sample is a linear function of sample-specific motif activities. Here, the motifs represent sets of TFs with shared binding specificity, i.e., non-redundant motif clusters as defined in HOCOMOCO v14^213^ and filtered, leaving only 585 motif clusters containing at least one transcription factor with CPM >0 in any CAGE-Seq muscle sample. Next, we run `maradoner create`, `fit`, `predict`, `grn` with `--no-means` and `export` in order to estimate mean and relative activities of 478 motif clusters passing MARADONER default filters for a minimal activity variance, see **Supplementary Data 8**, followed by testing their inter-muscular variation and contribution to the TRE expression. The ANOVA-like test for differential activity across muscles was performed via a Wald test performed for motif activity variances, see Section 8 in Meshcheryakov *et al.*^211^. Motif contribution to the TRE activity was estimated by computing the Bayesian belief, i.e., comparing the likelihoods under H_0_, the full model, against H_1_, constructed without the motif in question, see Section 10 in Meshcheryakov *et al.*^211^. To distinguish transcriptional activators and repressors, for each motif cluster we calculated the Pearson correlation coefficient between its non-standardized (raw) activity estimates for 222 muscle samples and the summary expression, log_2_ total CPM, of the corresponding transcription factors. For case studies, the high-scoring individual binding sites were identified with MoLoTool^213^ (https://molotool.autosome.org/).

### Allele-specific analysis

SNV calling, allele-specific read counting, and allelic imbalance estimation were performed using the approaches described in Viestra *et al.*^217^ and Buyan *et al.*^47^ and included the following steps:

1. Filtering out reads aligned with more than 2 mismatches and mapping quality (MAPQ) < 10 for all 225 BAM files generated in the previous step, along with the 3 files for non-muscle *septum intermusculare brachii laterale* samples, using the Python script filter_reads.py (https://github.com/StamLab/stampipes/tree/encode-release/scripts/bwa/, accessed in March 2025);
2. Variant calling using bcftools (v.1.19) mpileup with --redo-BAQ --adjust-MQ 50 --gap-frac 0.05 --max-depth 10000 and call with --keep-alts --multiallelic-caller;
3. Splitting the resulting SNVs into biallelic records using bcftools `norm` with `--check-ref x -m -` followed by filtering with bcftools `filter -i "QUAL > = 10 & FORMAT/GQ > = 20 & FORMAT/DP > = 10" --SnpGap 3 --IndelGap 10`, leaving only variants covered by 10 or more reads;
4. Annotating SNVs using bcftools `annotate` with `--columns ID,FREQ,COMMON`, and dbSNP (v.155), and leaving only biallelic sites using bcftools `view -m2 -M2 -v snps`;
5. Filtering variants with bcftools `query` and awk, leaving only those located on the autosomes with GQ ≥ 50, depth ≥ 10, and allelic counts ≥ 5;
6. Applying WASP (v.0.3.4) together with HISAT2 and `filter_reads.py` to find, remap, and filter the SNV-overlapping reads that failed to map back to the same location after the allele swapping according to the WASP procedure;
7. Running Python scripts count_tags_pileup_new.py and recode_vcf.py from https://github.com/autosome-ru/MixALime/tree/main/natcomm_supp_scripts, accessed in December 2025, to perform sample-level allelic read counting and BED to VCF conversion.

Finally, we applied MIXALIME (v.2.27.3) to 222 VCF files corresponding to muscle samples passing quality filters. Particularly, we ran mixalime `create --min-cnt 2`, `fit NB`, `test`, and `combine` by muscles with `--adaptive-min-cover`. Given the low number of biological replicates, from 2 to 4, we decided to use the negative binomial model. Finally, all ASVs were annotated with ANNOVAR (v.2020Jun7)^218^ `table_annovar.pl -buildver hg38 -protocol refGene -operation g` using RefSeq gene annotation downloaded from UCSC in February 2025.

### Partitioned heritability analysis

To perform heritability partitioning^179^ using ASVs obtained on the previous step, we selected GWAS statistics on 41 muscle-related phenotypes available at Pan UK Biobank (https://pan.ukbb.broadinstitute.org/) and 17 from NHGRI-EBI GWAS Catalog^178^, see **Supplementary Data 9**. Stratified linkage disequilibrium score regression analysis (S-LDSC, v.1.0.1) was performed as described on the LDSC partitioned heritability GitHub page (https://github.com/bulik/ldsc/wiki/Partitioned-Heritability); the necessary files, including the baseline-LD model (v.2.2), were downloaded from S-LDSC ZENODO (v.4)^219^. First, GWAS full summary statistics were lifted over to the hg38 genome with liftOver (v.447) and the UCSC chain file, followed by converting them to a .sumstats file using munge_sumstats.py script and HapMap3^220^ SNP list. Second, .annot files were created using bedmap from BEDOPS (v.2.4.41) and previously downloaded 1000G PLINK .bim files. Third, baseline and target annotation LD scores were calculated with `ldsc.py --l2 --ld-wind-cm 1.0` command. Finally, to partition heritability by functional categories `ldsc.py --h2 --overlap-annot --print-coefficients` was executed for each target annotation separately. For categorical traits, we additionally used `--samp-prev` and `--pop-prev`, setting them equal (for UKBB) or estimating phenotype population prevalence by literature searching (for potentially biased studies from NHGRI-EBI). S-LDSC results for different muscle ASV categories are available in **Supplementary Data 9**. Genetic correlations were estimated using `ldsc.py --rg` with the .sumstats files generated previously.

Following the filtering criteria from Finucane *et al.*^179^, we removed phenotypes with the total SNP-heritability z-score < 7 and left only one trait from a pair with a genetic correlation > 0.95, which resulted in 32 muscle-related phenotypes. Finally, Benjamini-Hochberg multiple testing correction for the enrichment *P*-values was performed separately for each ASV category, setting the `p.adjust = "BH"`.

### Validation of selected allele-specific variants

#### Plasmids

##### SNV reporter vectors

Promoters with a spliced 5’ UTRs of *MTOR* (mTOR), *RPL32*, *SYPL2*, and *ZDHHC5* were cloned into a promoterless reporter vector, pNL2.2 (Promega, Madison, WI, USA), by SLIC (sequence and ligation independent cloning) assembly, as described in Eliseeva *et al.*^221^. The length of the promoter region (see **Supplementary Data 10**) was determined based on the CAGE signal^48^, the ENCODE4^222^ registry of candidate Cis-Regulatory Elements (cCREs), and the presence of H3K27Ac mark. The promoters and 5’ UTRs were obtained from HEK293T genomic DNA, which was purified using GenElute Mammalian Genomic DNA Miniprep Kit (Merck KGaA, Darmstadt, Germany) according to the manufacturer’s instructions. The resulting plasmids were then sequenced to identify the first allele. The promoters containing the second allele were obtained from the corresponding plasmids using a four-primer PCR strategy and subsequently cloned into the same vector.

##### TF expression vectors

pcDNA-HA-ER Y537S^223^, expressing a constitutively activated form of ER (estrogen receptor), was a gift from Sarat Chandarlapaty (Addgene plasmid #49499; http://n2t.net/addgene:49499; RRID:Addgene_49499). The pcDNA3.1 HA-NRF1 plasmid was created by ligating the *NRF1* coding region, amplified from TFORF2523^224^ (gift from Feng Zhang, Addgene plasmid #142523; http://n2t.net/addgene:142523; RRID:Addgene_142523), into the *BamH*I and *Xho*I digested pcDNA3.1 HA-YY1 vector^225^ (gift from Richard Young, Addgene plasmid #104395 ; http://n2t.net/addgene:104395 ; RRID:Addgene_104395).

#### Cell cultivation and transfection

HEK293T cells (originally obtained from ATCC) were cultivated using a standard method in DMEM (Dulbecco Modified Eagle Medium) supplemented with 10% fetal bovine serum (Thermo Fisher Scientific), 2 mM glutamine, and 1x antibiotic-antimycotic. The cells were maintained at 37°C in a humidified atmosphere containing 5% CO_2_.

HEK293T cells were transfected using Lipofectamine 3000 (Thermo Fisher Scientific) according to the manufacturer’s protocol. 50 ng of pNL2.2 plasmid alone (for MTOR and SYPL2) or the mixture of 50 ng of pNL2.2 and 200 ng of appropriate pcDNA3.1-HA were used for each 48-well. After 16 hours, the cells were collected for RNA purification or for luciferase activity measurement.

### NanoLuc luciferase activity (NlucP) measurement

Cells were lysed in Passive Lysis Buffer (PLB) (Promega) for 10 min at 37°C, and luminescence was measured using the Nano-Glo Luciferase Assay System (Promega) and a GloMax 20/20 Luminometer (Promega).

#### RNA purification and quantitative real-time PCR (qPCR)

The total RNA was purified using QIAzol Lysis Reagent (Qiagen, Germany) and Direct-Zol RNA Microprep kit (Zymo Research, USA) according to the manufacturer’s recommendations. dsDNase treatment and reverse transcription were performed using the Maxima H Minus First Strand cDNA Synthesis Kit (Thermo Fisher Scientific). qPCR was performed on a DTlite Real-Time PCR System (DNA Technology) using qPCRmix-HS SYBR+LowROX reaction mixture (Evrogen) and NlucP-specific primers (**Supplementary Data 10**) according to the manufacturer’s recommendations. The two-tailed Student’s t-test was conducted to estimate the statistical significance between the reference and alternative alleles.

### Summary of the external annotations used in the study

#### Annotation of FANTOMUS TREs

The following external genomic annotations were used to characterize 35895 autosomal FANTOMUS TREs.

1. FANTOMUS annotation by chromatin states was performed using bedtools intersect with the three groups of BED files. First, we downloaded 6 BED files from ENCODE^52^ containing all available histone modification ChIP-Seq peaks for the reference *m. gastrocnemius caput mediale* epigenome (ENCSR108YWO). Second, all 12 narrowPeak files from Roadmap Epigenomics^51^ containing histone ChIP-Seq and DNase-Seq peaks for HSMM cell-derived skeletal muscle myotubes (E121) were downloaded and converted to hg38. Third, 5 BED files corresponding to CUT&Tag peaks obtained from *m. vastus lateralis* for different histone modifications were downloaded from GEO under the accession number GSE195854^53^.
2. Comparison to FANTOM5^48^ was performed for 203052 and 61825 autosomal promoters and enhancers downloaded from the FANTOM hg38 collection (fair+new peaks, https://fantom.gsc.riken.jp/5/). Only the intersections with a maximum overlapping fraction of 0.5 or more were considered, i.e., TREs intersecting by ≥ 50% of either FANTOMUS or FANTOM5 cluster length. We also excluded intersections of the FANTOMUS unidirectional TREs with the FANTOM5 promoters located on the opposite strands. Muscle promoters were defined as the regions with > 0.5 CPM in both adult skeletal muscle samples (FANTOM5 IDs 10023-101D5 and 10282-104F3) or all 4 EOM (10272-104E2, 10297-104G9, 10298-104H1, and 10299-104H2). For muscle enhancers, we additionally required their usage according to F5.hg38.enhancers.expression.usage.matrix in either both somitic or all extraocular muscles. This resulted in 33948 and 90 muscle promoters and enhancers, respectively. Eye-specific FANTOM5 promoters were defined as the regions with > 0.5 CPM in all 4 eye muscle samples and neither of the 2 skeletal muscles, which resulted in 591 TREs. Additionally, we downloaded Supplementary Table 4 from the FANTOM5 article^48^ to annotate a subset of 10077 intersected FANTOM5 TREs as house-keeping promoters.
3. We also performed the TRE-level comparison against the CAGE-Seq data obtained from the biopsy samples of *m. vastus lateralis* and *m. soleus* from Makhnovskii *et al.*^55^ and Bokov *et al.*^193^, respectively. The former included 43527 autosomal TSS clusters downloaded from GEO under the accession number GSE164081, which were compared to FANTOMUS TREs in the same way as FANTOM5. For the latter, we obtained 8 FASTQ files from the authors corresponding to 4 individuals before and after dry immersion; for comparison, we repeated the procedure described in the "CAGE-Seq data processing" section, followed by calculating median log_2_CPM.

#### Genes and proteins annotation

To perform gene-level comparison with GTEx (v.10, RNASeQCv2.4.2)^54^, we used median gastrocnemius muscle transcripts per million (TPM) estimates for 35384 autosomal protein-coding and lncRNA genes. Of them, 16860 were also detected in FANTOMUS and used for estimating total gene-level correlation, and 13405 with gastrocnemius median TPM > 0.5 were used for **Figure 1e**.

Proteins identified with MS were divided into functional groups using a list of 171 ribosomal proteins obtained in July 2025 from HGNC^226^ (HGNC Group ID 1054), 135 translation-associated proteins from Anisimova *et al.*^227^, 1615 transcription factors from HOCOMOCO v14^213^ and 236 proteins involved in muscle contraction retrieved from AmiGO2 (isa_partof_closure: GO:0006936, taxon_subset_closure_label: Homo sapiens, aspect: P, type: protein) which were not annotated by the previous protein sets.

#### Muscles annotation

The classification of 27 leg muscles into three groups based on their susceptibility to 25 myopathies was performed by manual curation of the available literature listed in **Supplementary Data 5**. Groups were defined as usually spared (0), affected later (1), and affected earlier and severely (2). For gene expression ANOVA, we considered 18 causative genes corresponding to myopathies with at least 5 different muscles in each group.

#### Annotating ASVs

FANTOMUS ASV annotation was performed using 1621950 skeletal muscle eQTLs from the GTEx (v.10)^54^ Muscle_Skeletal.v10.eQTLs.signif_pairs.parquet file, 190162 ADASTRA (v.6.1)^176^ ASBs significant (FDR < 0.05) for at least 1 transcription factor, 11020 UDACHA (v.1.0.3)^47^ ASVs significantly (FDR < 0.05) affecting chromatin accessibility in 49 muscle-related samples, and 17386 SNVs from 18818 conditionally distinct eQTL signals from Wilson E.P. *et al.*^177^.

## Data availability

The complete FANTOMUS data, including TRE- and gene-level counts, protein group abundances, MIXALIME allelic imbalance estimates, and MARADONER output tables, are available on Zenodo^228^ (https://zenodo.org/records/19389795). The raw CAGE-Seq reads are deposited to GEO under accession number GSE310878. The mass spectrometry proteomics data have been deposited to the ProteomeXchange Consortium via the PRIDE^229^ partner repository with the dataset identifier PXD072528.

## Code availability

The source code of the CAGE-Seq peak calling pipeline is available on GitHub (https://github.com/autosome-ru/fantomus-clustering).

## Author contributions

**Writing - Original Draft, Writing - Review & Editing:** I.V.K., A.B., V.M., I.A.E., A.R.R.F., D.P., E.S.

**Data Curation, Investigation, Methodology:** I.V.K., A.B., G.G., E.S., V.G.Z., N.E.V., I.A.E., Y.S., A.T., L.S., A.M., R.D.

**Formal Analysis, Software, Investigation, Methodology:** I.V.K., A.B., N.G., V.N., A.V.E., G.M., D.P.

**Conceptualization:** Y.H., A.R.R.F., O.G., E.S., D.P., I.V.K.

**Project Administration, Funding Acquisition, Resources, Supervision:** I.V.K., V.M., D.P., E.S., Y.H., O.G.

## Acknowledgments

The mass spectrometry-based proteomics analysis was performed on the equipment of the "Human Proteome" Core Facility in the Institute of Biomedical Chemistry (Moscow). We thank Ekaterina Kulakovskaya for the schematic depiction of human skeletal muscles.

## Figure Legends

**Supplementary Figure 1. Evaluating CAGE-Seq data.**

**a**, Total number of reads (top), percentage of reads mapped to the reference genome (middle), and number of uniquely mapped reads (bottom) for 228 samples. Color denotes body part-based muscle group, data range, and means are indicated in the middle. **b**, The results of bulk CAGE-Seq data deconvolution for 225 skeletal muscle samples. Samples were hierarchically clustered based on cell type fractions using the default complete linkage method and Euclidean distances; color denotes cell type fraction. Left: muscle group colored as in (**a**) and quality status (failing samples are black). **c**, PCA of the uncorrected gene-level log_2_CPM values for 225 skeletal muscle samples. Axes: the first two components, color denotes individuals, shape reflects the sample quality status, with three filtered samples labeled. **d**, Reference bias, defined as the fraction of variants with the reference allelic read coverage exceeding the alternative allele, for 225 skeletal muscle samples. X-axis: the total number of reads covering SNV; color: muscle group; shape: sample quality status. **e**, Categorical distribution of uni- (top) and bidirectional (bottom) TREs across genomic regions. **f**, Fractions of uni- and bidirectional TREs intersected with ChIP-Seq peaks for histone modifications from the Roadmap Epigenomics HSMM (top), ENCODE skeletal muscles (bottom left), and Galle *et al.* (bottom right). CAGE-Seq: cap analysis of gene expression, PCA: principal component analysis, MF: myofiber, MSC: muscle stem cell, SMC: smooth muscle cell, EC: endothelial cell, FB: fibroblast, Ref: reference allele, TRE: transcribed regulatory element, CDS: coding sequence, UTR: untranslated region, CPM: counts per million, SNV: single-nucleotide variant, HSMM: human skeletal muscle myoblasts.

**Supplementary Figure 2. Calling CAGE-Seq peaks to identify FANTOMUS TREs.**

**a**, Pipeline used for peak calling, see **Methods**. Briefly, CAGE-Seq 5’-profiles were used to identify CTSS (1), divide their coverage into soft-clipped and non-clipped (2) and filter out positions where these metrics can not be applied (3), call bidirectional TREs (4), annotate them and filter out promoter regions (5), call unidirectional TREs (6), annotate all TREs together (7) and perform filtration by total and soft-clipped read coverage (8-9). The final set of TREs used in FANTOMUS was obtained by intersecting the result with MANE and GENCODE CDS to remove the signal likely originating from the "exon painting". **b**, The distribution of the softclip score, log_10_ ratio of the number of soft-clipped reads to the number of non-clipped, for 1360769 TREs from step (8) in (**a**). Panels represent different genomic regions; dashed line: threshold used for TRE filtering. TRE: transcribed regulatory element, biTREs: bidirectional TREs, uniTREs: unidirectional TREs, CAGE-Seq: cap analysis of gene expression, CTSS: CAGE transcription start site, softG: reads with soft-clipped 5’-G, noclip: reads without soft-clipped 5’-end, CPM: counts per million, CDS: coding sequence, UTR: untranslated region.

**Supplementary Figure 3. FANTOMUS CAGE-Seq comparison to external datasets and MS.**

**a**, Correlation between median log_2_TPM from GTEx (X-axis) and log_2_CPM from FANTOMUS (Y-axis) for 16860 genes; solid line represents diagonal. **b**, The expression level of 345 FANTOMUS TREs intersecting eye-specific FANTOM5 promoters. Boxplot whiskers demonstrate the minimum and the maximum, except for outliers with values further than 1.5 interquartile range from the hinge; boxes reflect first, second, and third quartiles, and color denotes muscle group. *P*-value: two-tailed paired Student’s t-test for TRE log_2_ mean CPM in EOM against all other muscles. **c**, Top: distribution of the softclip score, log_10_ ratio of the number of soft-clipped reads to the number of non-clipped, for 1307091 autosomal verifiable CAGE clusters, both passing and failing FANTOMUS quality filters. Linetype: cluster quality status; color: intersection with FANTOM5. Middle: log_2_ median CPM; boxplots are built as in (**b**); color denotes intersection with FANTOM5. Bottom: categorical distribution of TREs of different quality status across genomic regions. Bars are filled and colored by gene and FANTOM5 annotations, respectively. **e**, Basic MS quality metrics: total number of detected peptides (top) and number of peptides per protein (bottom) for the 4 batches, i.e., sets of isobaric labels, each containing 15 samples and one control (all samples pooled together, see **Methods**). **f**, The percentage of genes detected in MS (red) and CAGE-Seq (purple) that belong to different biological groups; the total number of detected genes is indicated to the right. **g**, GSEA results for genes ordered by the residuals from **Figure 1h**. Only 34 gene sets demonstrating FDR < 0.05 and |NES| > 2 are shown. **h**, Violins demonstrating the sample-level SCC between transcript (log_2_CPM from CAGE-Seq) and protein (log_2_ reporter ion intensity from MS) abundances. CPM: counts per million, TPM: transcripts per million, PCC: Pearson correlation coefficient, CDS: coding sequence, UTR: untranslated region, CAGE-Seq: cap analysis of gene expression, MS: mass spectrometry, GSEA: gene set enrichment analysis, NES: normalized enrichment score, SCC: Spearman correlation coefficient.

**Supplementary Figure 4. Muscle heterogeneity at the transcriptome and proteome levels.**

**a**, GSEA results for the EOM, tongue, and diaphragm differential gene expression and protein abundance analyses; genes and proteins were ordered by log_2_FC of the corresponding muscle against all the others. Only the top-10 and top-5 significantly enriched GO biological process pathways are shown for the transcriptome- and proteome-based results, respectively. Color denotes muscle group. **b**, GSEA results (left) for genes (right) ordered by their contribution to the first (top) and second (bottom) PCs of the transcriptome-based PCA, **Figure 2c**. Only pathways passing FDR < 0.05 and NES > 1.2 are left; the top 20 genes for each component are depicted on the right. **c**, GSEA results (left) for proteins (right) illustrated in the same way as in (**b**). **d**, Examples of the HOX gene expression patterns across the human body. Color denotes gene-level log_2_CPM. **e**, PC1 loadings from transcriptome- (X-axis) and proteome-based (Y-axis) PCA as in **Figure 2d**. Genes encoding proteasome subunits (left), complex I subunits (middle), and ribosomal proteins (right) are highlighted. **f**, Scatter plot demonstrating the skewness (X-axis) and bimodality coefficient (Y-axis) for 1541 proteins from 1469 protein groups significantly differentially abundant across muscles, ANOVA FDR < 0.05. Individual examples discussed in the text are highlighted and labeled; shape and color denote the expression pattern. **g**, Schematic representation of, left-to-right, *SIM1*, *EMX2*, *METTL21C* and *METTL21EP* expression distribution over human skeletal muscles; color scale reflects log_2_CPM. GSEA: gene set enrichment analysis, NES: normalized enrichment score, GO: gene ontology, PCA: principal component analysis, CPM: counts per million, EOM: extraocular muscles.

**Supplementary Figure 5. Examples of genes and proteins with differential expression and abundance across skeletal muscles.**

**a**, The transcription level (log_2_CPM) and protein abundance (log_2_ reporter ion intensity) distribution for the selected stretch-sensing genes; top: gene symbol and FDR-corrected ANOVA *P*-value; color denotes muscle group. **b-d**, The transcription level and protein abundance of the genes related to muscle atrophy (**b**), involved in muscle development and regeneration (**c**), and associated with muscular dystrophy (**d**), are shown as in (**a**). CAGE-Seq: cap analysis of gene expression, MS: mass spectrometry, CPM: counts per million.

**Supplementary Figure 6. Differential TRE usage across skeletal muscles.**

**a**, Top: distribution of log_2_FC calculated for 21235 TREs as their maximum log_2_CPM across muscle - maximum log_2_CPM of other TREs within the same gene, see **Methods**. Bottom: log_2_FC (X-axis) and maximal log_2_CPM across muscles (Y-axis) of the same TRE set. Dashed lines: |log_2_FC| = 5. **b**, Schematic representation of *ALPK2* short and long isoform expression difference distribution across muscles, color corresponds to log_2_FC. **c**, Genomic view (top), mean CAGE-Seq profiles for EOM, tongue, diaphragm, upper and lower limbs (middle), and FANTOMUS TRE track for the corresponding strand (bottom) for, top-to-bottom: *PITX2*, *SEPTIN9,* and *LIMS1*. Color denotes muscle group. FDR for EOM, tongue, and diaphragm was obtained by performing differential TRE usage analysis with edgeR. **d**, The transcription level (log_2_CPM) and protein abundance (log_2_ reporter ion intensity) distribution for the selected TREs and protein isoforms; top: ANOVA *P*-values, FDR-corrected for TRE differential activity; color denotes muscle group. CAGE-Seq: cap analysis of gene expression, MS: mass spectrometry, FC: fold change, CPM: counts per million, FDR: false discovery rate.

**Supplementary Figure 7. Differential motif activity across muscles.**

**a**, PCA of the 75 skeletal muscles based on the activities of 353 motif clusters passing ANOVA FDR < 0.05. Axes: the first two components; color denotes muscle group. **b**, Schematic representation of the expression distribution of, left-to-right then top-to-bottom: MEF2 target *CSRP3* promoter, HSF target *HSPA6*, THR targets *HR* and *MYLK4*, ALX target *MYH15* and ZBTB7A target *ATP1A3*; color scale reflects log_2_CPM. **c**, PCA based on the gene-level log_2_CPM values of 42 fast- and slow-twitch marker genes as in **Figure 2f**. Axes: the first two components; color: motif activity. **d**, The transcription level (log_2_CPM) distribution for the selected progenitor cell markers; top: gene symbol and FDR-corrected ANOVA *P*-value; color denotes muscle group. CAGE-Seq: cap analysis of gene expression, PCA: principal component analysis, CPM: counts per million.

**Supplementary Figure 8. Allele-specific variants affecting muscle TRE activity.**

**a**, Percentage of muscle ASVs annotated as, top-to-bottom: skeletal muscle eQTLs from GTEx, ASBs from ADASTRA, ASVs affecting muscle chromatin accessibility from UDACHA, and eQTLs from Wilson E.P. *et al.*; X-axis: allelic imbalance FDR cutoff. Solid pink lines: percentages for individual muscles; dashed black line: an average trend; vertical line: FDR = 0.05. **b**, Stratified LD score regression results for muscle ASVs passing certain allelic imbalance significance thresholds and 32 GWAS traits. Color scale reflects regression coefficient; size and stroke color denote the enrichment and FDR, respectively. GWAS traits were hierarchically clustered with cosine distances and UPGMA using pairwise genetic correlations. eQTL: expression quantitative trait loci, ASB: allele-specific transcription factor binding site, ASV: allele-specific variant, LD: linkage disequilibrium, GWAS: genome-wide association study, FDR: false discovery rate, UPGMA: unweighted pair group method with arithmetic mean.

