## Supplementary Figures for "A Joint Promoterome-Proteome Atlas Highlights the Molecular Diversity of Human Skeletal Muscles"

Supplemetary Figure 1

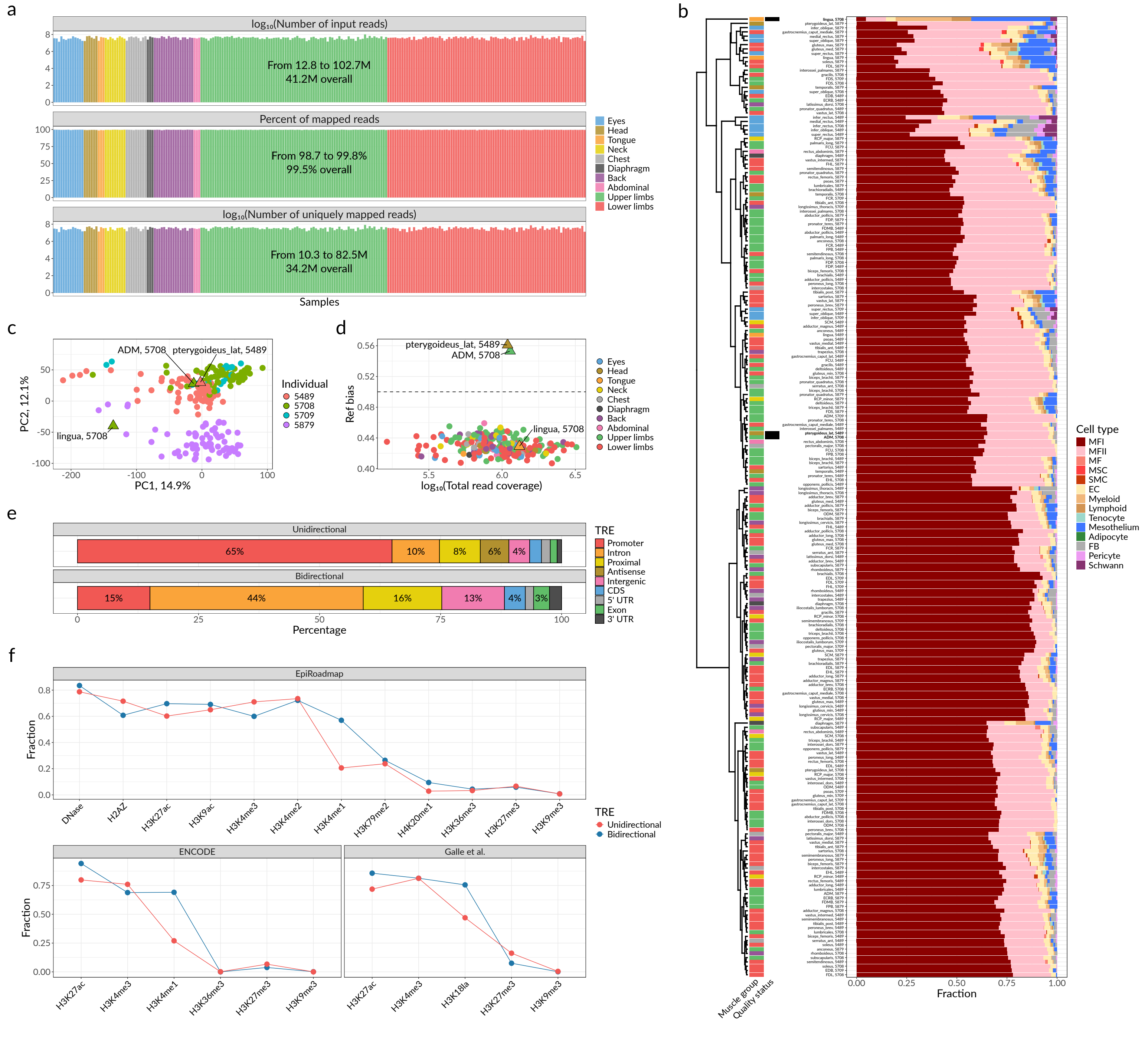

Supplemetary Figure 2

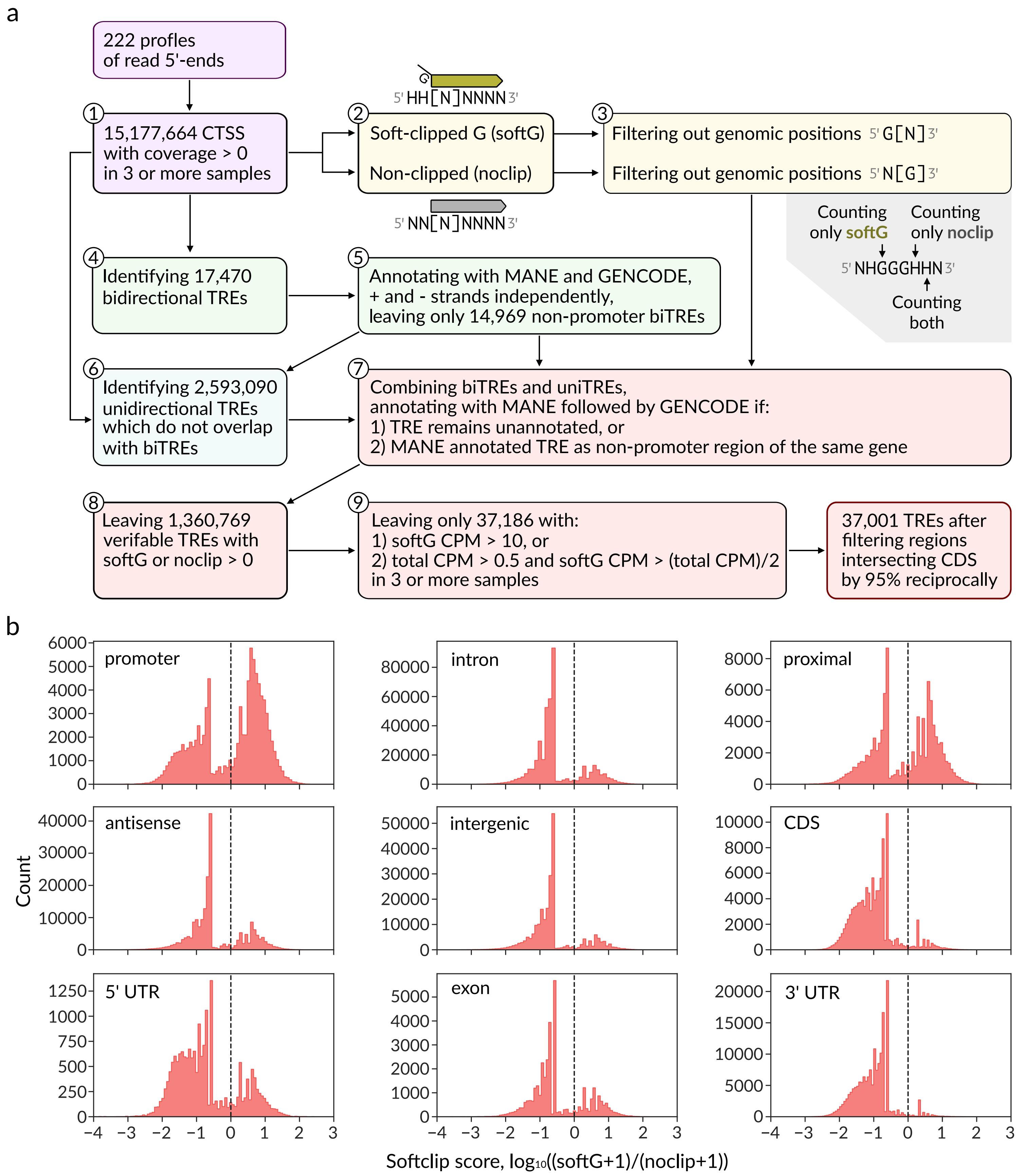

Supplemetary Figure 3

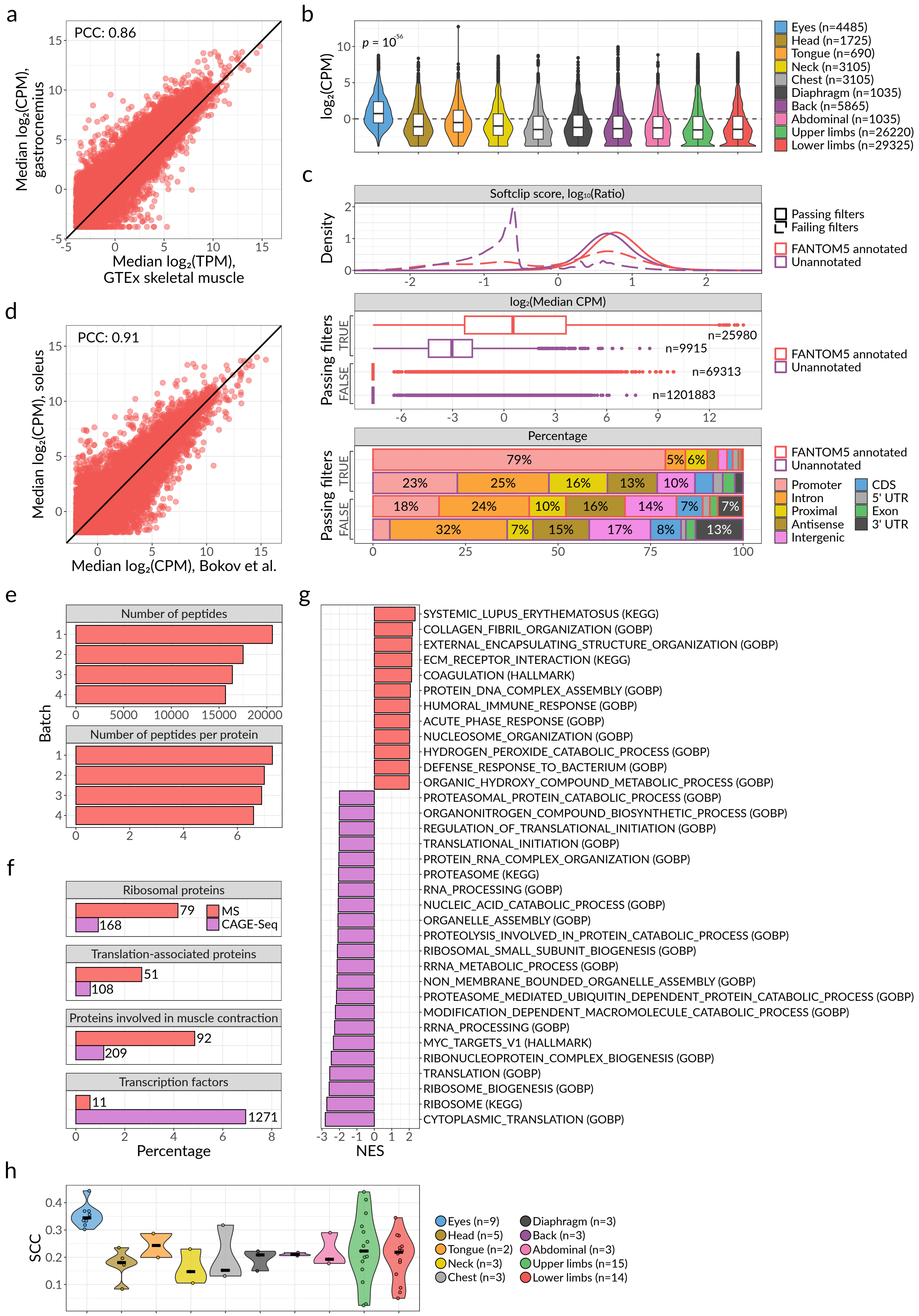

Supplemetary Figure 4

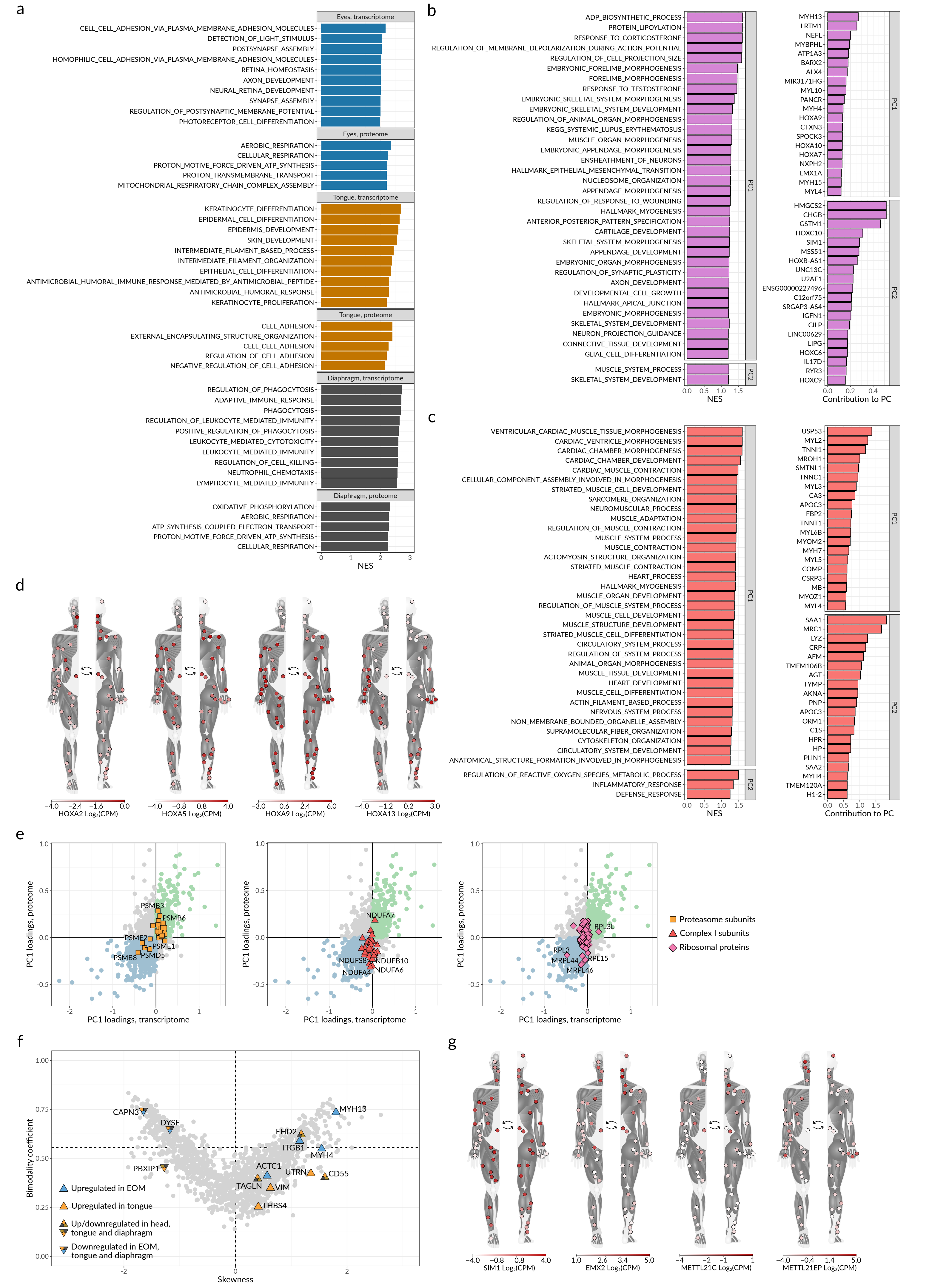

Supplemetary Figure 5

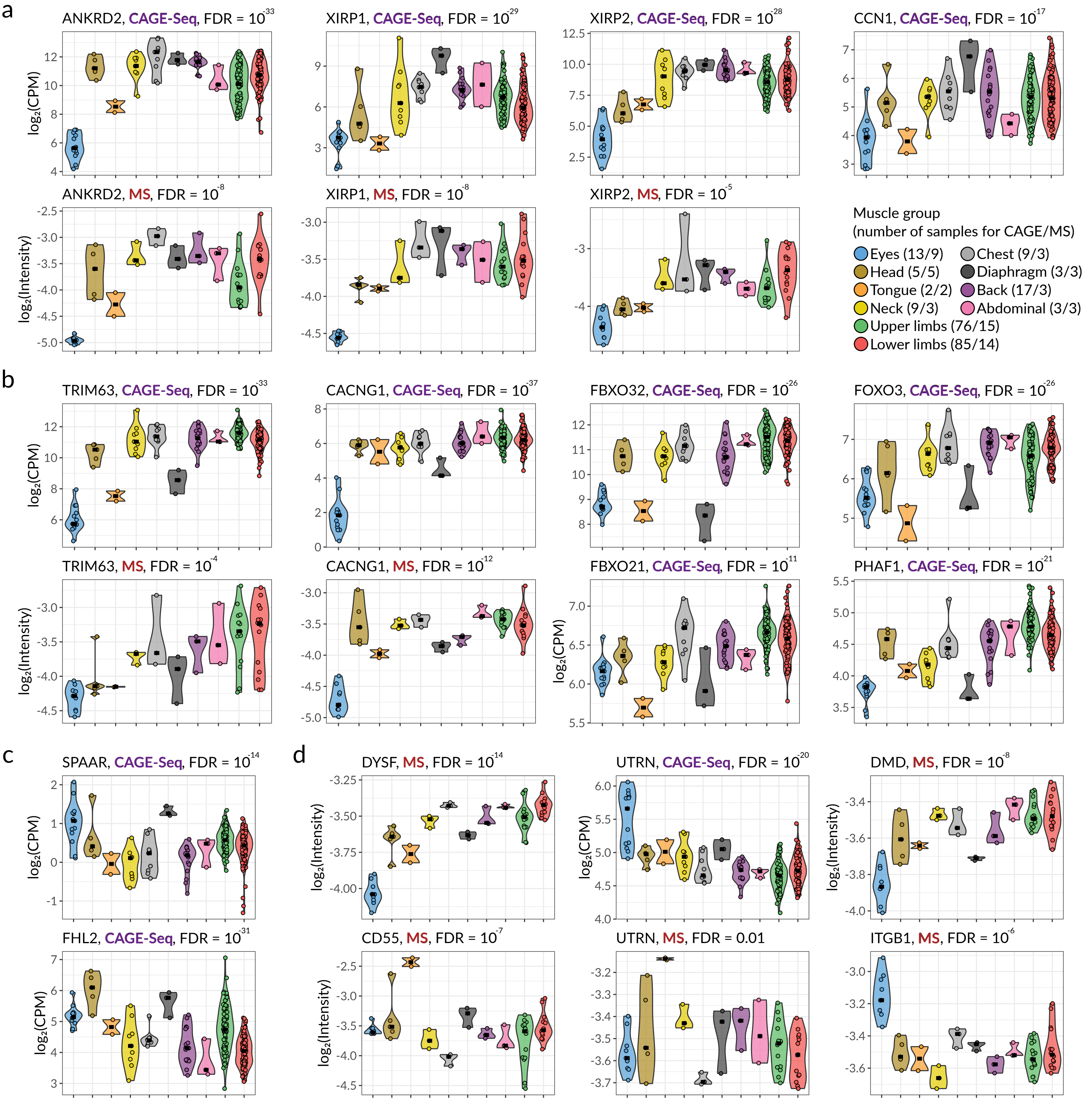

Supplemetary Figure 6

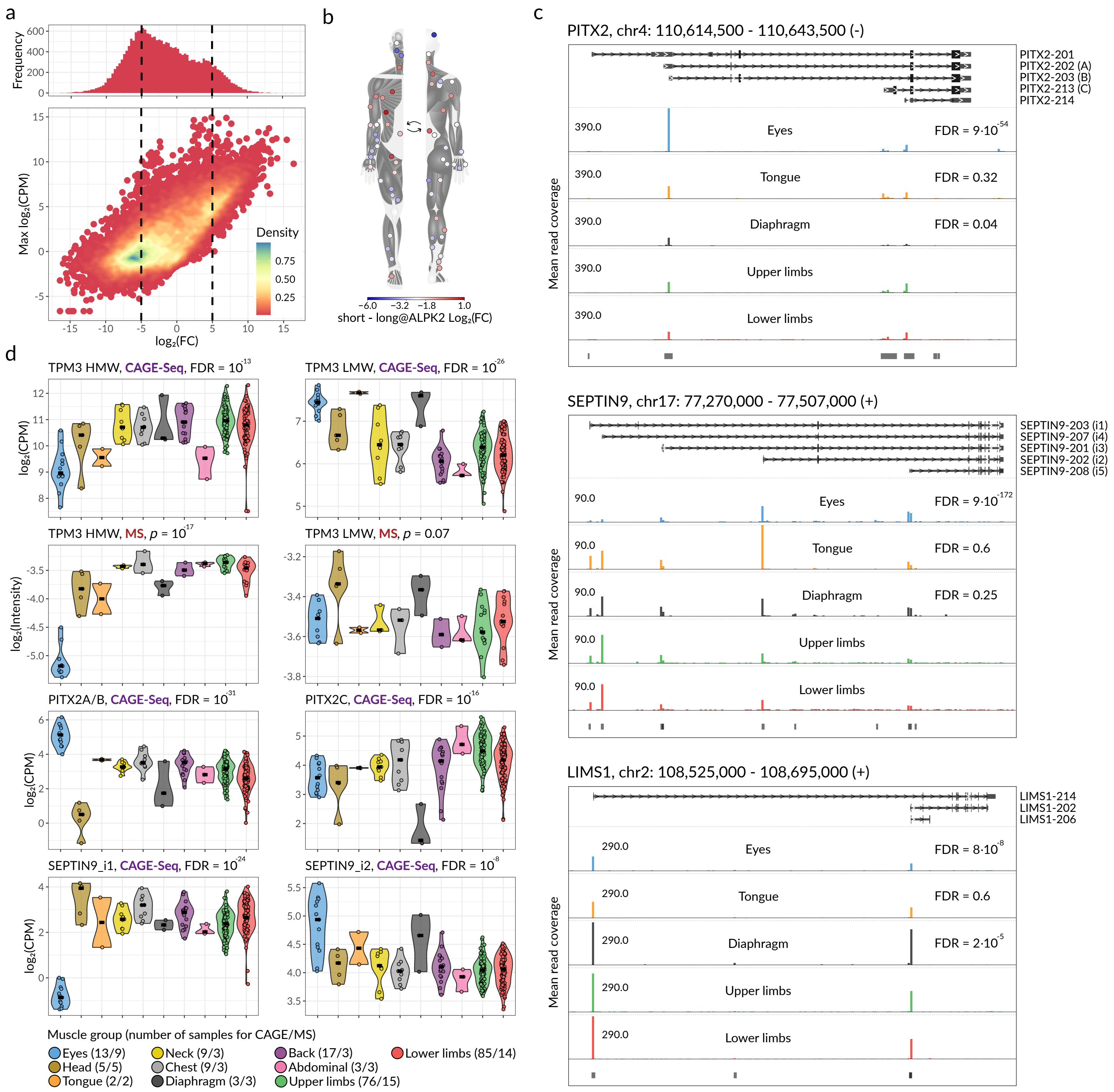

Supplemetary Figure 7

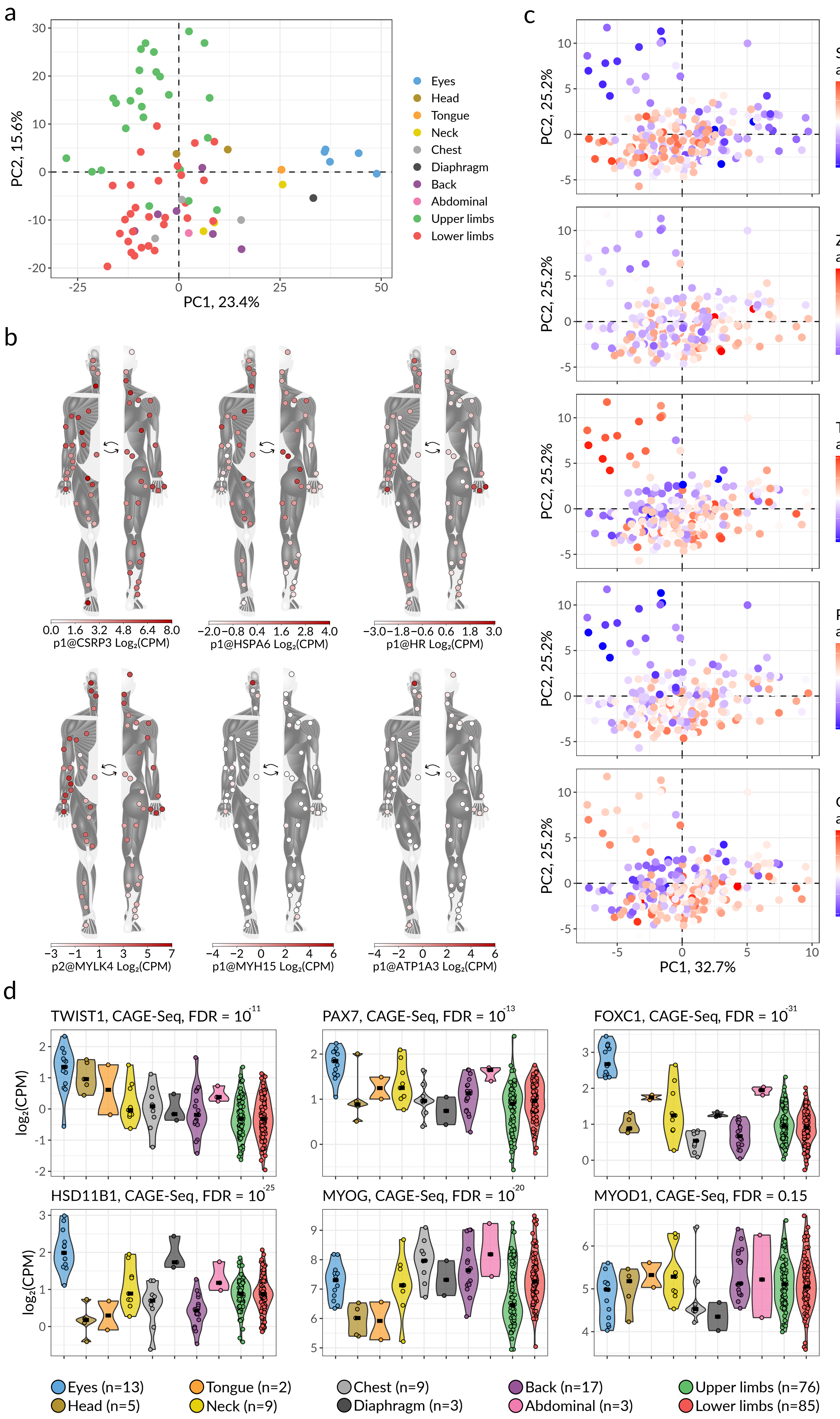

Supplemetary Figure 8

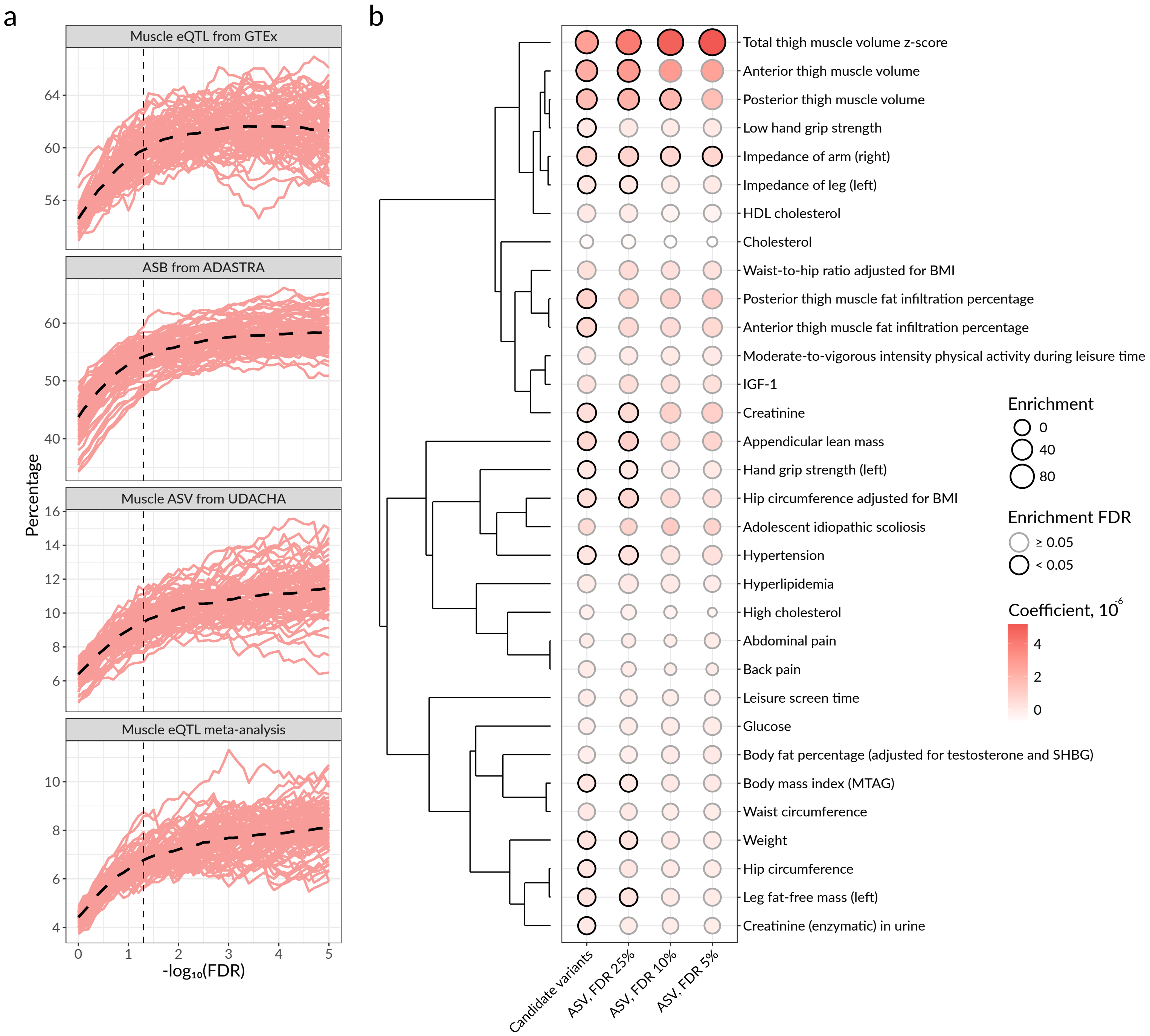
